# Systematic Optimization Enables Near-Perfect In Vitro Transformation Efficiencies for *Spirodela polyrhiza* (Greater Duckweed)

**DOI:** 10.1101/2025.09.10.675371

**Authors:** Tasmia Islam, Ayalew Ligaba-Osena, Eric A. Josephs

## Abstract

The in vitro transformation of plants, or the delivery of foreign genetic material that is incorporated into their genomes, represents a powerful tool both for elucidating genotype-phenotype relationships and for generating plant cultivars which have desirable traits for agriculture and/or biotechnological applications. However, outside of a few model species, the processes involved in transformation are often inefficient and can take months to perform for many plant species, with several bottlenecks occurring at the different stages of calli induction, genetic transfection, and plant regeneration. While duckweeds – aquatic monocots whose species include some of the smallest and fastest-growing flowering plants on the planet – have distinguished themselves with several emerging biotechnological applications, they too are the subject of conflicting reports regarding their transformation potential and ability to be genetically manipulated. Here, we synthesized and optimized the protocols for in vitro transformation of duckweed *Spirodela polyrhiza* (Greater Duckweed) from start-to-finish: achieving >90% - 100% efficiencies for each of calli induction; transient and stable genetic transformation; visual marker-free selection of transformants; and regeneration of genetically modified plants with stable transgene expression for over 100 generations – and which in *S. polyrhiza* can be achieved over the course of weeks instead of months. The integrated, streamlined approaches for all stages of in vitro transformation overcome many bottlenecks and can help to pave the way for high-throughput functional genomics studies and synthetic biology applications in this biotechnologically-important species.

## Introduction

Duckweeds (*Lemnaceae*) represent a family of free-floating, monocotyledonous aquatic plants that are widely distributed across freshwater habitats worldwide (1). Comprising five genera—*Spirodela*, *Landoltia*, *Lemna*, *Wolffiella*, and *Wolffia*—duckweeds exhibit highly reduced morphologies, consisting primarily of a frond or thallus, and reproduce predominantly through vegetative budding from meristematic pockets (1,2). Historically, *Spirodela polyrhiza* has gained prominence as a model for studying developmental processes such as leaf heterophylly (3) and plant senescence (4), owing to its short life cycle (5), simple structure, and genetic accessibility (6,7). Advances in transcriptomics and genome editing have deepened our understanding of key developmental and physiological processes, including metabolic regulation, leaf morphogenesis, and circadian rhythm control (8–13). Beyond their utility in basic plant biology, duckweeds have attracted considerable attention for environmental applications, particularly in ecotoxicology and phytoremediation, due to their exceptional capacity for bioaccumulation and contaminant uptake (14,15). Additionally, duckweeds’ high starch and protein content, coupled with their ability to thrive in nutrient-rich wastewater, makes them attractive candidates for biofuel production, animal feedstocks, and other biorefinery applications (16–19).

With their small genomes; rapid vegetative propagation where daughter fronds bud from a mother plant every <30 hours (and biomass doubling approximately every 2 day); ease of cultivation under controlled conditions; and demonstrated capacity for stable transgene expression—such as in *Spirodela oligorrhiza*, where GFP expression exceeds 25% of total soluble protein, placing it among the highest expressing nuclear transformation systems in higher plants (20–22) – duckweeds have emerged as promising platforms for biotechnology and synthetic biology. Among duckweed species, *S. polyrhiza* is distinguished by its relatively large frond size (up to 1.5 cm) and the smallest genome within the family, estimated at 158 Mb (23). The publication of the *S. polyrhiza* draft genome by Wang et al. (2014), followed by a high-resolution chromosomal assembly by Michael et al. (2017), and more recently, the genome sequences of five duckweed species by Martienssen et al. (2025), have collectively laid the groundwork for genomic and biotechnological investigations. Their simplicity makes them ideal for high-throughput genetic studies, functional genomics, and metabolic engineering.

Because their simplicity makes them ideal for high-throughput functional genomics studies and applications of plant synthetic biology and metabolic engineering, high-efficiency in vitro transformation (delivery and incorporation of foreign genetic material into their genomes) can be especially impactful in species like duckweeds (Figure 1). However, outside of a few model plant species such as *Arabidopsis thaliana* and *Nicotiana benthamiana*, many plant species can be particularly recalcitrant to both the delivery and incorporation of foreign genetic material into their genomes and the regeneration of transfected tissue back into full plants. Often, these processes are time consuming and labor-intensive, requiring significant, month-long efforts with only small rates of success; even in well-studied plants such as soybean, sorghum, or switch-grass, transformation efficiencies often are <10% (24). In particular, many plants are recalcitrant to infection by *Agrobacterium* species to deliver transgenes to plants, which can be highly dependent both on the plant cultivar and the strain of *Agrobacterium*, with success rates are often <1 - 2% for transgene delivery. For their part, genetic engineering strategies have been successfully demonstrated in duckweeds, enabling the production of recombinant proteins, biodegradable polymers, and high-value metabolites for pharmaceutical and industrial purposes (25–29). Callus induction (30–32), nanomaterial-mediated plasmid transfection (9), genetic transformation (31,33), and regeneration (30,31) systems have been successfully established for *S. polyrhiza*, each with variations of protocols and components of growth media with reported efficiencies. Similar transformation protocols have also been developed for other duckweed species, including *Spirodela punctata* (34), *Lemna gibba* (35), *Lemna minor* (35), *Wolffia arrhizal* (36), and *Wolffia columbiana* (37)*, Wolffia globosa* (38) highlighting the growing interest in duckweeds as promising platforms for plant biotechnology. However, even when these processes are optimized individually, it is often found that the success rates of each of these processes can be dependent on one other, such that efficient transformation requires successful end-to-end optimization of tissue culture, callus induction, transformation, and plant regeneration to be efficient.

**Figure 1.**
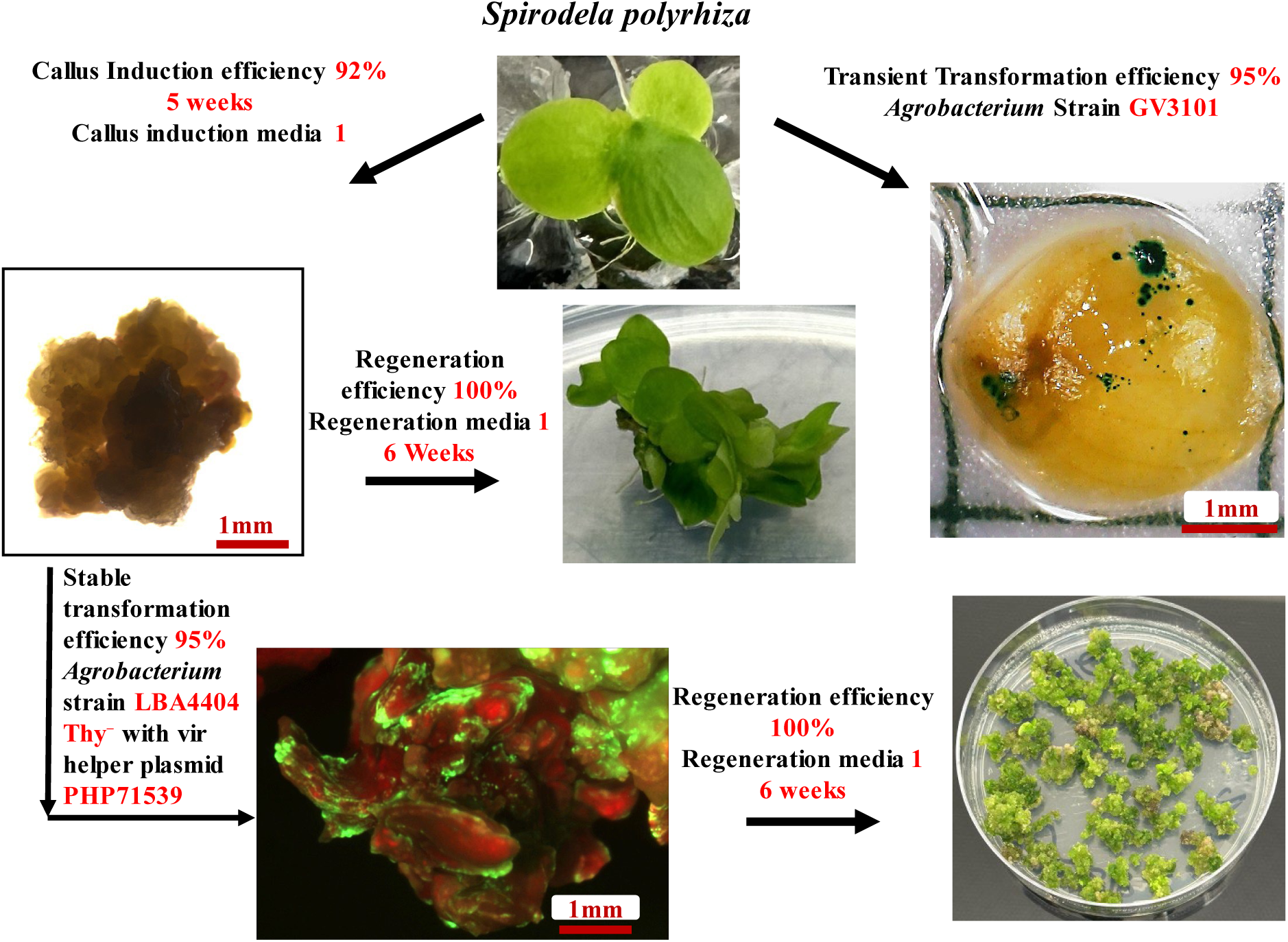
“Bottleneck-free” in vitro transformation of greater duckweed (*Spirodela polyrhiza*) from end-to-end. In this report, conditions for tissue culture; calli induction; transient and stable transformation by *A. tumefaciens*; selection and plant regeneration are reported with each step taking a few weeks and achieving >90 – 95% efficiencies at each stage for *S. polyrhiza*.

Given the increasing interest in *S. polyrhiza* for genetic engineering and bio-production applications, here we performed a systematic, comparative analysis across the multiple published protocols, as well as variations thereof, to optimize callus induction, genetic transformation (*Agrobacterium tumefaciens*-mediated transient and stable), clonal selection, and regeneration as a high-efficiency, integrated platform for *S. polyrhiza* synthetic biology. Highlighting explant age as a key determinant of success, callus induction rates as high as 92% could be achieved, followed by the identification of specific combinations of callus induction media, explant age, and regeneration conditions that could lead to regeneration efficiencies of up to 100%. At the same time, we also were able to identify the most effective strains of *A. tumefaciens* for both transient and stable transformation of transgenes, reaching 90% - 95% efficiencies for transient transformation, and for the first time evaluating the effects of maize morphogenic regulator genes *Baby boom (Bbm)* and *Wushel (Wus2)* on improving the efficiency of stable transformation in duckweeds (39). Because of the streamlined protocols, over 100 calli were generated and tested for each transformation condition, allowing us to demonstrate 95% stable transformation efficiencies, including the delivery and visual marker-free selection of the genes for CRISPR-Cas9 that are used for precision genome editing in plants. Regenerated plants expressing transgenes could be maintained for over 270 days (∼115 generations). By our synthesizing the divergent protocols for *S. polyrhiza*, *S. polyrhiza* is situated as a high-throughput platform for its myriad emerging applications in functional genomics, synthetic biology, and plant biotechnology that is optimized for ease and extreme efficiency: as fast-growing monocot that rapidly generates clonal “daughter” plants vegetatively, made easy-to-transform with near-perfect efficiencies in just weeks, we expect duckweeds such as *S. polyrhiza* can emerge as the “*E. coli*” of plant synthetic biology.

## Results

### Optimization of Media for Callus induction in Duckweed

Mature, fully expanded mother fronds aged 1 to 2 weeks (Figure 2A), with rhizoids removed or shortened, were manually separated from younger daughter fronds and used as explants for callus induction. These mature fronds were chopped and plated (∼20 explants per plate; Figure 2B) onto seven different callus induction media (CIM) formulations (Table 1), then incubated in the dark at 28°C for five weeks. Callus induction efficiency was assessed by recording the number of explants that formed calli (Figures 2C-D). A significant variation in callus induction efficiency was observed among the seven media tested (Figure 2E). CIM1 produced the highest induction rate of 92% using explants from 2-week-old cultures. CIM2 and CIM3 also showed intermediate induction, each achieving approximately ∼72%, while CIM4 through CIM6 yielded 63%, 47%, and 20%, respectively. No callus formation was observed on CIM7. Overall, explants derived from 2-week-old cultures exhibited significantly higher callus formation compared to those from 1-week-old cultures, indicating that explant physiological maturity strongly influences morphogenic competence. After five weeks, the calli were transferred to fresh CIM1 for an additional one week of proliferation. Morphological progression of callus development— from fronds to explants to callus—is illustrated in Figures 2A–D, confirming successful induction (Figure S1) and proliferation (Figure S2).

**Figure 2.**
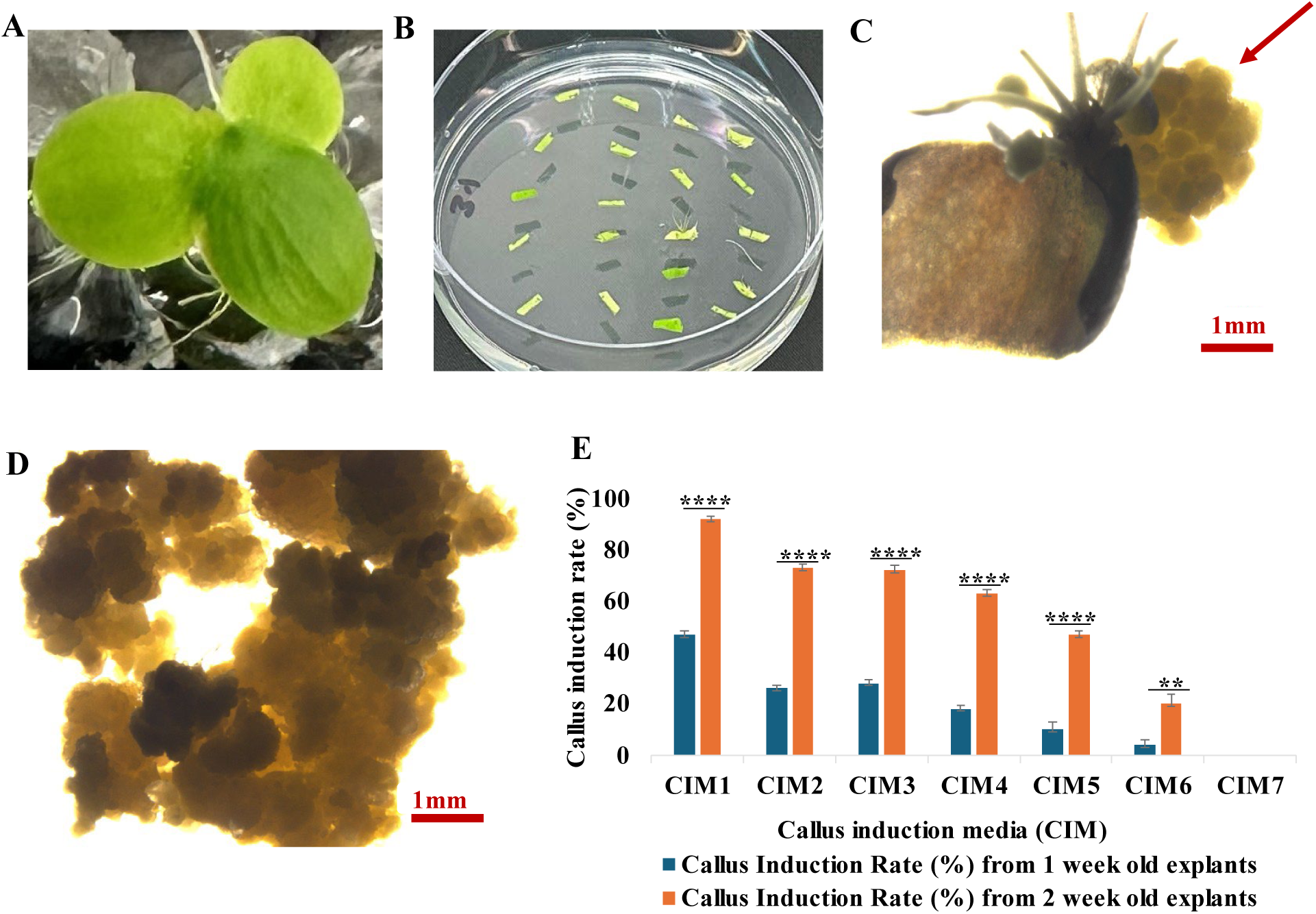
Callus induction in *S. polyrhiza* on CIM. (A) Intact fronds. Fronds were prepared as described in Materials and Methods. (B) Chopped explants plated on CIM. (C) Callus initiation (red arrow) from explant tissue. (D) Proliferating friable callus clusters. Scale bars = 1mm. (E) Callus induction efficiency was evaluated after 5 weeks on seven CIM formulations. CIM1 gave the highest induction rate (92%) with 2-week-old explants, whereas no callus was obtained on CIM7. Values represent mean ± SE (n = 5). Statistical differences between explant ages were determined by Student’s *t*-test (**p < 0.001; ****p < 0.0001).

**Table 1.**
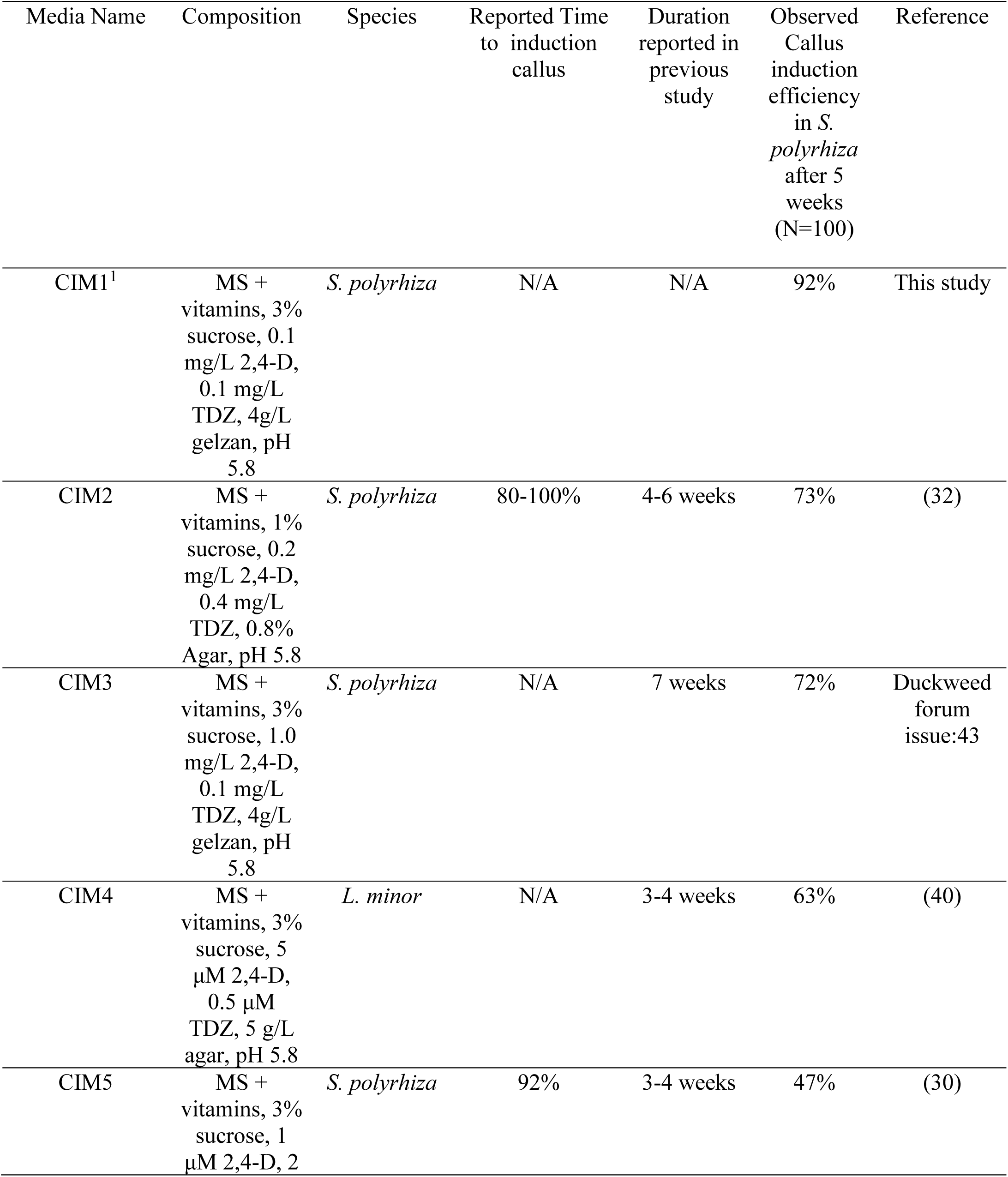
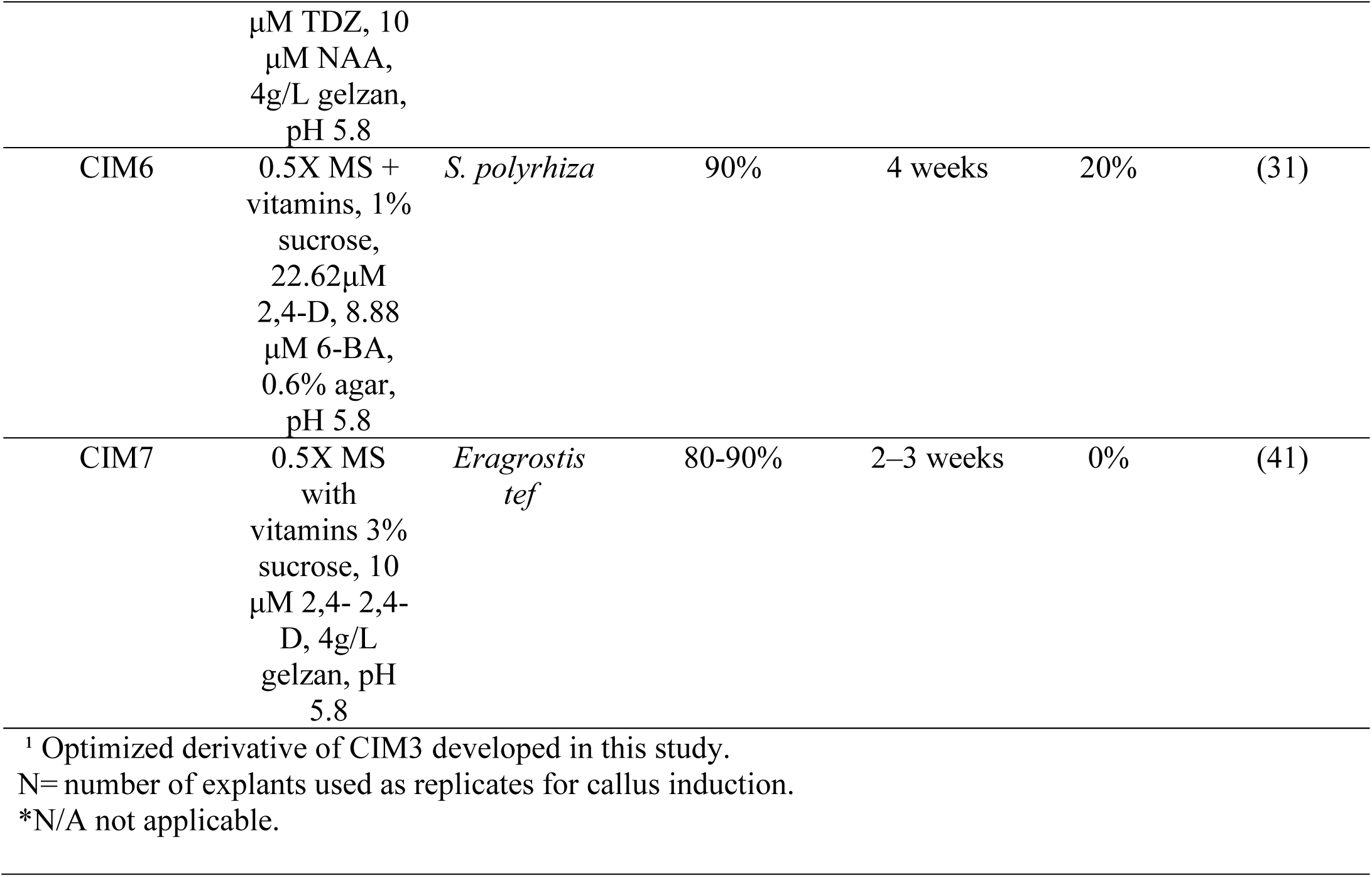
Conditions for Callus Induction Evaluated.

**Table 2.** Conditions for Regeneration Efficiency Evaluated.

| Media Name | Composition | Species | Reported<br>regeneration<br>efficiency in<br>previous study | Duration<br>reported for<br>regeneration<br>in previous<br>study | Observed<br>regeneration<br>efficiency in<br>*S. polyrhiza*<br>within 6<br>weeks<br>(N=50) | Reference |
| --- | --- | --- | --- | --- | --- | --- |
| RM1 | 0.5X MS media,<br>1% (m/v) sucrose,<br>0.6% (m/v) agar,<br>4.54 µM TDZ, pH<br>5.8 | *S. polyrhiza* | 96% | N/A | 100% | (31) |
| RM2 | SH media, 10 µM<br>zeatin, 1% sucrose,<br>0.6% (m/v) agar,<br>pH 5.8 | *S. polyrhiza* | 70-80% | 10 weeks | ≥90% | (32) |

N= number of calli used as replicates for regeneration.
\*N/A not applicable.

### Optimization of Media for Callus Regeneration in Duckweed

Calli induced on each of the seven callus induction media (CIM) were subsequently transferred to two regeneration media, RM1 and RM2, and evaluated for their ability to regenerate into whole *S. polyrhiza* plants under 16h/8h day/night cycles at 28°C. After transfer from dark to light, the beige calli began turning green within 2–3 days on both RM1 and RM2 media (Figures 3A–B), followed by maturation of the calli (Figure 3C). New frond regeneration from these embryonic calli was observed after approximately 3–4 weeks (Figures 3D and S3). By 6 weeks, all calli had fully transformed into clusters of fronds (Figure 3E). These clusters were then transferred to liquid 0.5X SH medium, where they developed rhizoids and disaggregated into individual fronds that could be easily separated to form independent plants (Figure 3F). On RM1, all calli showed 100% regeneration efficiency regardless of the callus induction media used (Figure 3G). In contrast, regeneration efficiency on RM2 varied depending on the CIM: calli derived from CIM1, CIM2, and CIM3 exhibited the highest regeneration rates (97.5%), followed by CIM4 (95%), CIM5 (90%), and CIM6 (82.5%) (Figure 3G).

**Figure 3.**
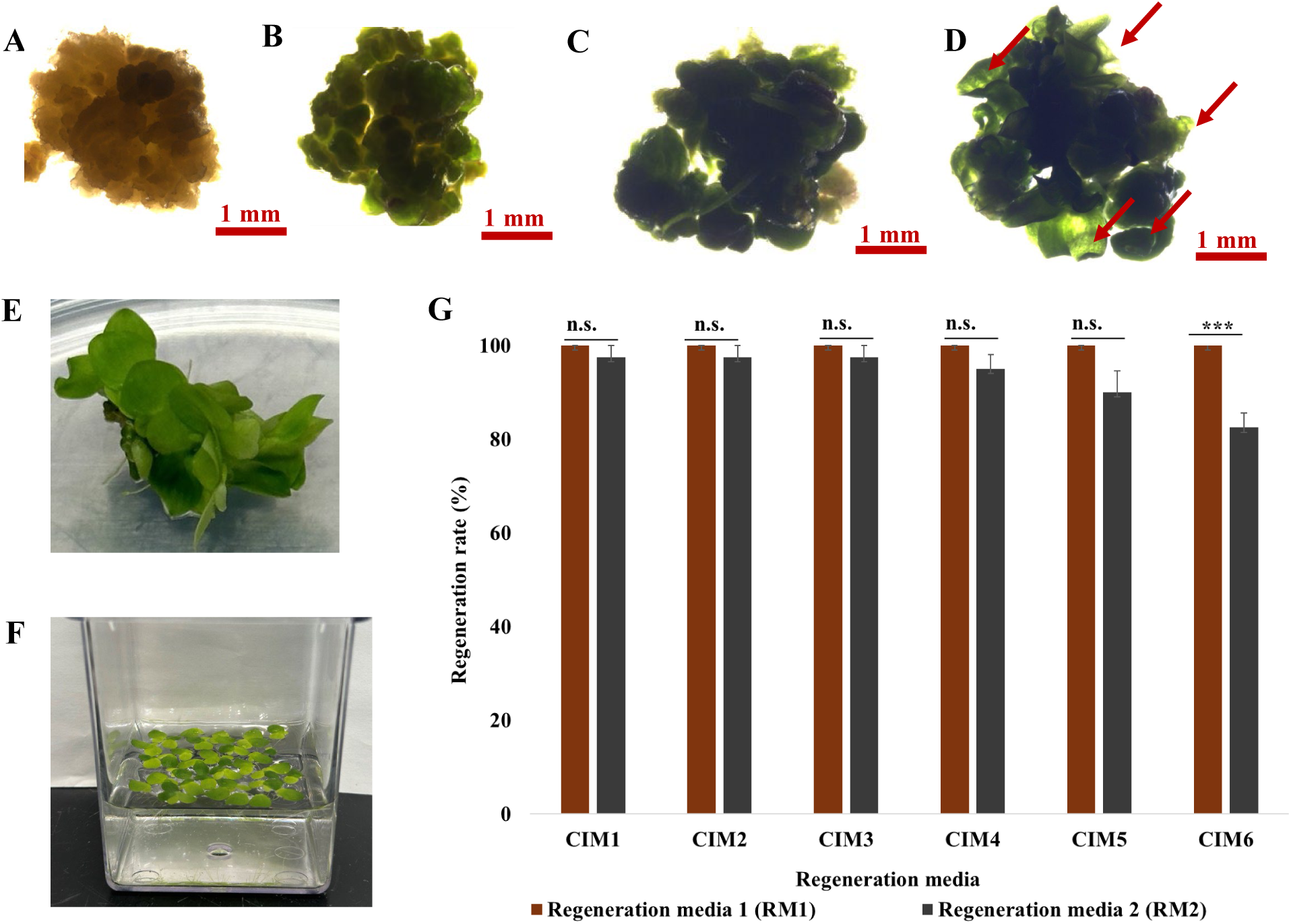
**Shoot regeneration and plantlet development from callus tissue of *S. polyrhiza.*** (A) Callus tissue on regeneration media. (B) Early chlorophyll accumulation visible as greening of callus tissue, followed by (C) maturation. (D) Development of shoot primordia and multiple emerging shoots from callus tissue (red arrows). (E) Fully developed plantlets with multiple fronds regenerated from callus. (F) Regenerated plantlets grown in vitro in liquid culture (0.5× SH medium with 1% sucrose, pH 5.8). Scale bars = 1 mm (A-D), 50 µm (E, F). (G) Regeneration efficiency of calli derived from different callus induction media on two regeneration media. RM1 supported 100% regeneration across all calli. On RM2, calli from CIM1, CIM2 and CIM3 showed the highest regeneration (97.5%), followed by CIM4 (95%) and CIM5 (90%), with CIM6 showing the lowest (82.5%). Data are presented as mean ± standard error (n = 5). Statistical significance was determined using Student’s t-test; “ns” denotes no significant difference, and *** indicates p < 0.001.

### *Agrobacterium* Mediated Transient and Stable Transformation

#### Optimization Transient Transformation of *S. polyrhiza*

We aimed to establish robust protocols for both stable and transient transformation in *S. polyrhiza* by evaluating two commonly used *A. tumefaciens* strains, GV3101 and LBA4404, for their ability to mediate gene transfer in *S. polyrhiza* (Table 3). GV3101, derived from the C58 background and harboring the disarmed Ti plasmid pMP90, is known for its high virulence and strong performance in dicot transformation, including *Arabidopsis* and duckweed species. LBA4404, derived from the Ach5 background and containing the disarmed pAL4404 plasmid, has historically been used in dicots and monocot transformations but often shows lower transformation efficiency in many plant species, particularly, monocots. Agrobacterium-mediated transformation is known to be attenuated by an immune response from the plant and many plants are recalcitrant to transformation by these means; however, duckweeds are known to have divergent immune-response genes compared to terrestrial plants which may allow for higher transformation efficiencies under proper conditions.(42)

**Table 3.** Transformation efficiency evaluation of different *Agrobacterium* strains.

| *Agrobacterium*<br>Strain | Trial | Total Fronds | Positive Fronds | Transformation<br>Efficiency (TE)<br>% | Average<br>TE<br>% |
| --- | --- | --- | --- | --- | --- |
| GV3101 | 1 | 25 | 23 | 92 | 95 |
|  | 2 |  | 24 | 96 |  |
|  | 3 |  | 24 | 96 |  |
| LBA4404 | 1 | 25 | 0 | 0 | 0 |
|  | 2 |  | 0 | 0 |  |
|  | 3 |  | 0 | 0 |  |

**Table 4.** Composition of Media Used in Stable Transformation.

| Media type | Composition |
| --- | --- |
| Co-cultivation media | 0.5X MS media, 4.58 mM MES, 22.62 µM 2,4-D, 8.88 µM 6-BA, 100 µM AS, 50 mg/L thymidine, pH 5.2 |
| Resting media | 0.5X MS media with vitamins, 1% sucrose, 0.6% agar, 9.99 mM (NH<sub>4</sub>)<sub>2</sub>SO<sub>4</sub>, 22.62 µM 2,4-D, 8.88 µM 6-BA, 0.63 mM cefotaxime, pH 5.8 |
| Selection media | For PHP81858: 0.5X MS media, 1% sucrose, 0.6% agar, 9.99 mM (NH<sub>4</sub>)<sub>2</sub>SO<sub>4</sub>, 22.62 µM 2,4-D, 8.88 µM 6-BA, 0.63 mM cefotaxime, 30 mg/L G418 (pH 5.8).<br>For JD633: 0.5X MS media with the same components but supplemented with 5 mg/L hygromycin instead of G418. |
| Excision media | 0.5X MS media, 1% sucrose, 0.6% agar, 9.99 mM (NH<sub>4</sub>)<sub>2</sub>SO<sub>4</sub>, 22.62 µM 2,4-D, 8.88 µM 6-BA, 50 µM ABA, pH 5.8 |
| Frond induction media | 0.5X MS, 1% sucrose, 0.6% agar, 4.54 µM TDZ, pH 5.8 |

The two agrobacterium strains were evaluated based on their delivery of plasmid pSB161, a binary plant transformation vector derived from the pCAMBIA backbone, designed for *A.*-mediated gene delivery. It carries a CaMV 35S promoter-driven GUS reporter gene and a kanamycin resistance gene *neomycin transferase* (nptII) for selection in plants. The vector is suitable for both transient and stable transformation in dicot species and is compatible with commonly used *Agrobacterium* strains such as GV3101 and LBA4404. Transient expression was confirmed through GUS histochemical assays, indicating successful expression of the reporter gene in transformed tissues (Figures 4B and S4). No blue regions were observed in control sample without agrobacterium (Figure S5).

**Figure 4.**
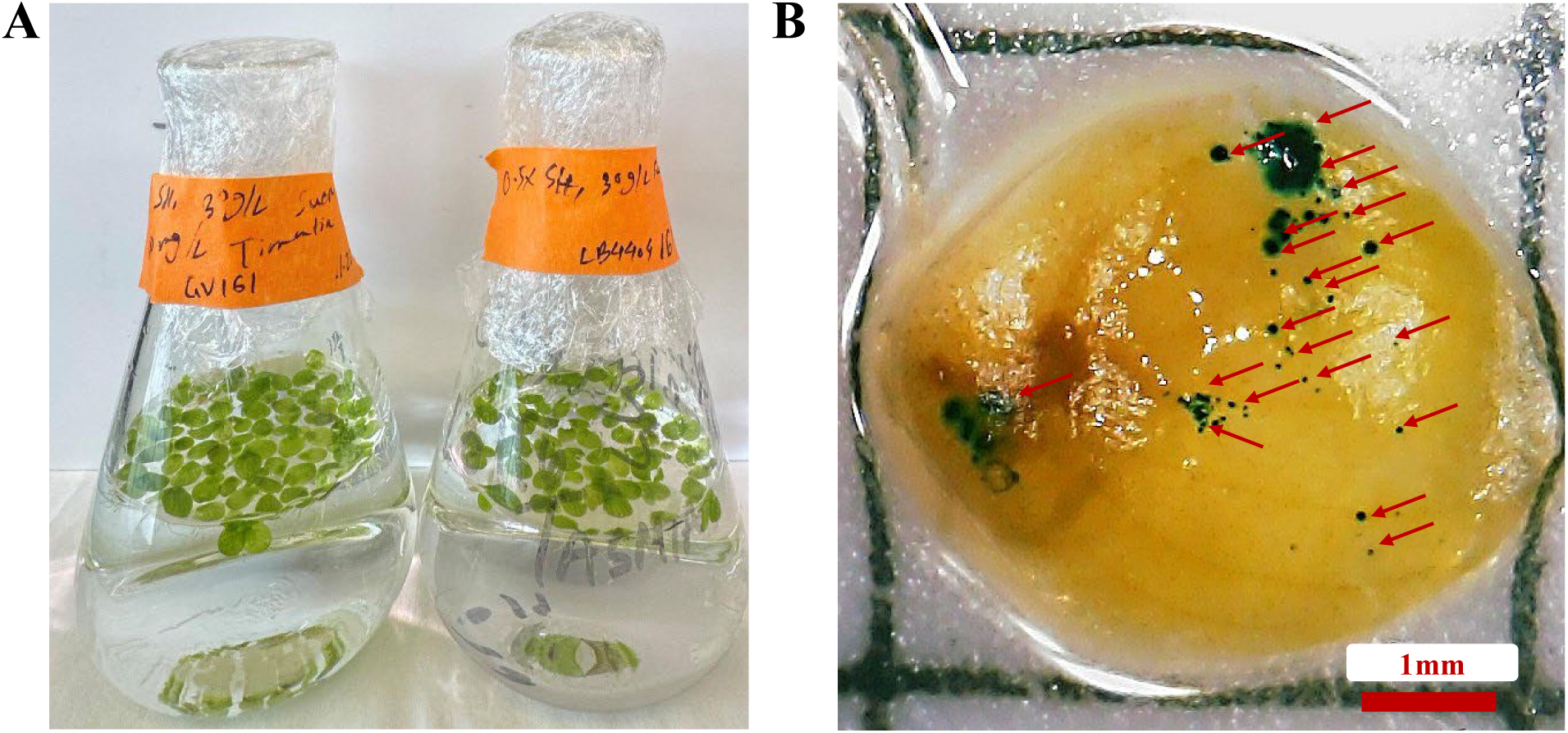
**Transient transformation of *S. polyrhiza* using *Agrobacterium tumefaciens.*** (A) Transformed fronds were cultured in liquid medium following co-cultivation. Only fronds treated with *A. tumefaciens* strain GV3101 exhibited signs of successful transient transformation, whereas those treated with LBA4404 showed no detectable response. (B) Histochemical GUS staining of *S. polyrhiza* callus tissue transformed with GV3101 harboring the pSB161 plasmid. Multiple, blue-stained foci (indicated by red arrows) represent regions of GUS reporter gene expression, confirming successful transient transformation and efficiency is 95%.

Optimization efforts included testing a range of variables, among which three factors emerged as critical for successful transformation: (i) vacuum infiltration, (ii) a 30-minute post-infiltration resting period, and (iii) the inclusion of 200 μM AS in both inoculation and co-cultivation media under dark conditions. Omission of any of these elements consistently led to failed transformation events. Following co-cultivation, positive GUS activity was observed in plants cultured in liquid medium only (Figure 4A). Our findings indicate that GV3101 outperforms LBA4404 in transient transformation assays using reporter construct such as 35S::GUS. Notably, transient transformation was not successful with strain LBA4404, which yielded no detectable transformants.

#### Stable Transformation of *S. polyrhiza*

To optimize the stable transformation protocol for *S. polyrhiza*, we conducted multiple trials using the auxotrophic *Agrobacterium* strain LBA4404 Thy⁻ carrying the virulence helper plasmid PHP71539 and the binary vector PHP81858 containing maize morphogenic regulator genes *Baby boom (Bbm)* and *Wushel (Wus2)* and AmCyan1 which is bright cyan fluorescent protein, obtained from Corteva Agriscience (39,43). This agrobacterium strain has been optimized for monocot transformation. It is defective in thymidine biosynthesis and requires thymidine supplementation in the media for growth.

Because calli induction was so efficient, this allowed us to generate up to 100 calli per condition in order to optimize and evaluate transformation efficiencies under those conditions (Table 5). Embryogenic calli were successfully transformed through a combination of shaking and vacuum infiltration, followed by co-cultivation on medium supplemented with acetosyringone and thymidine to promote T-DNA transfer. Optimal transformation was achieved using *Agrobacterium* cultures with an OD₆₀₀ of 0.8–0.9 and a 5-day co-cultivation period (Figures 5B–C). Post co-cultivation, explants were transferred to selection medium containing 30 mg/L geneticin (G418) for three weeks (Figure 5E). Bright GFP expression was detected in calli within the first week of selection (Figures 6A, S6), while no fluorescence was observed in non-transformed controls (Figure S7), confirming successful transgene delivery.

**Figure 5.**
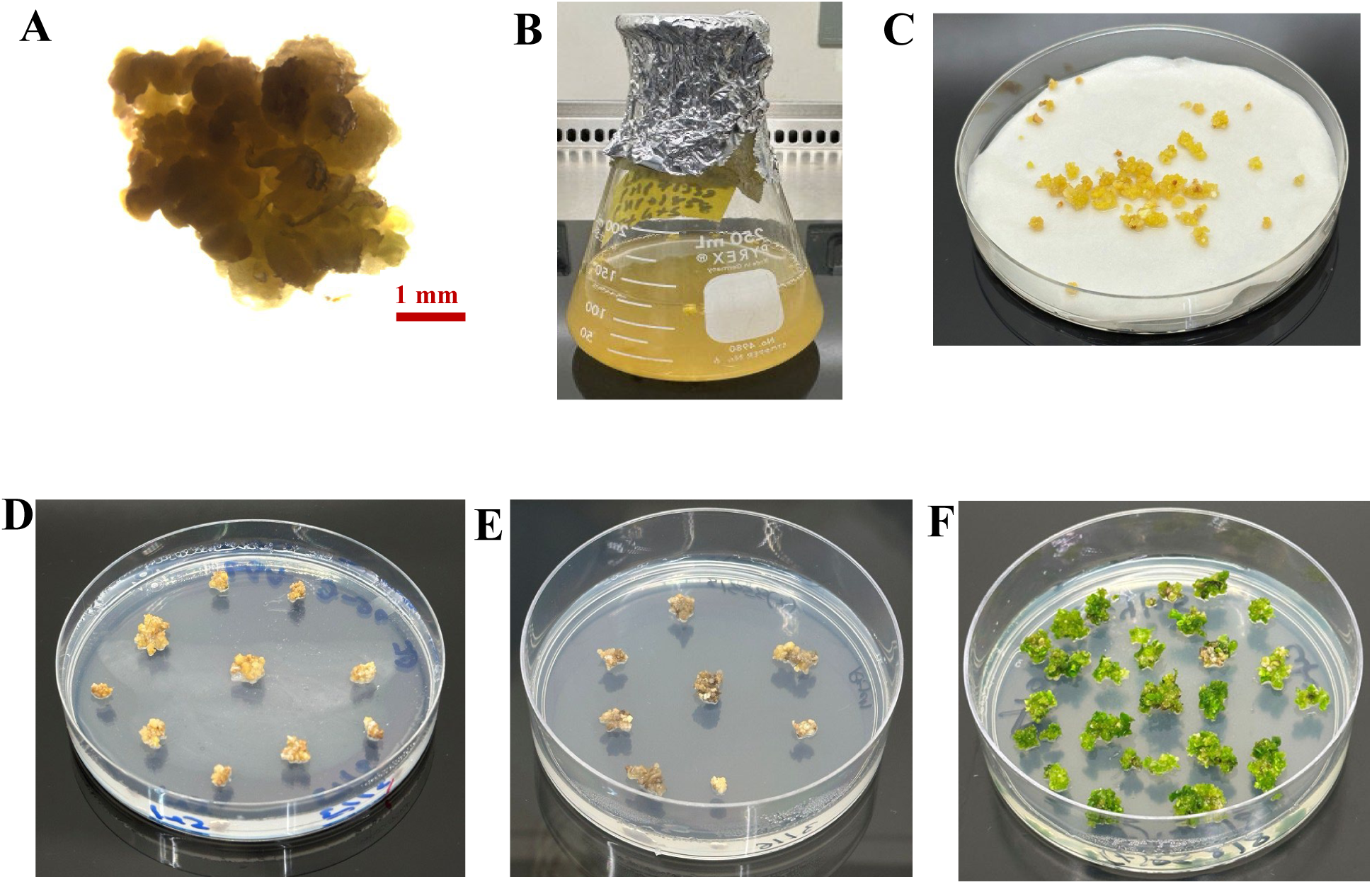
Stable and Efficient Genetic Transformation of *S. polyrhiza.* (A) Representative image of healthy callus tissue prior to transformation. (B) Co-cultivation of calli with *A. tumefaciens* in liquid suspension. (C) Transfer of infected calli to filter paper soaked in liquid co-cultivation medium. (D) Post co-cultivation, calli are placed on regeneration medium to suppress bacterial overgrowth and promote tissue recovery. (E) Selection of transgenic lines on medium containing 30 mg/L G418. (F) Emergence of transgenic fronds from selected calli (red arrows), showing greening and shoot-formation indicative of successful regeneration and transformation.

**Figure 6.**
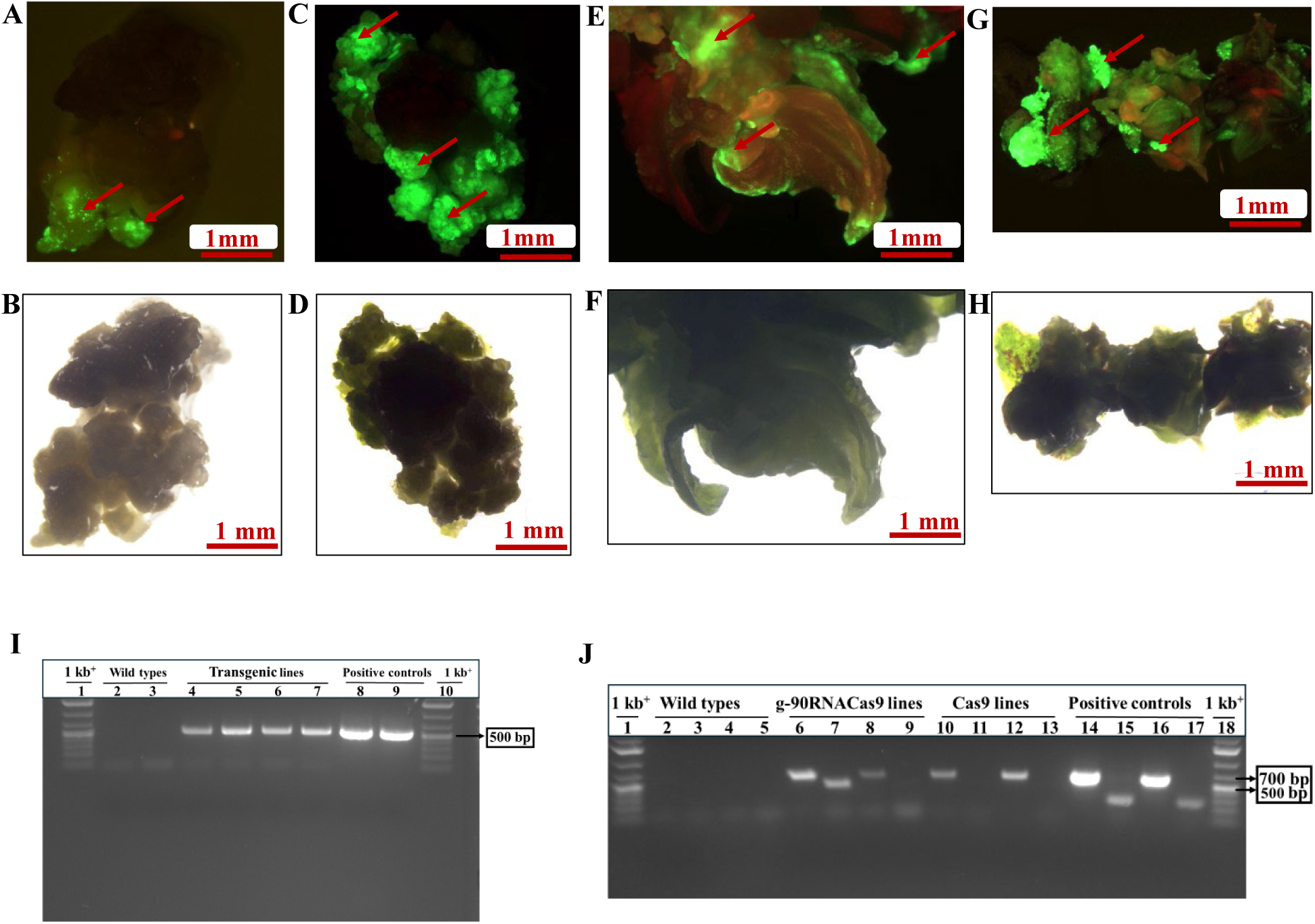
Stable transformation, regeneration, and molecular confirmation of S. polyrhiza transformed with GFP reporter, Cas9 and g-90RNACas9 constructs. (A) Callus tissue under G418 selection showing GFP fluorescence after transformation with *A. tumefaciens* LBA4404 Thy⁻ carrying the PHP81858 plasmid, indicating stable integration and expression of the GFP gene. (B) Corresponding bright-field image of the same callus tissue without fluorescence filter. (C, E, G) Regenerated fronds at 2, 6, and 9 weeks on regeneration media, exhibiting strong GFP fluorescence (red arrows), confirming sustained transgene expression during regeneration. (D, F, H) Corresponding bright-field images of regenerated fronds. (I) PCR confirmation of GFP integration in transgenic *S. polyrhiza*. Agarose gel electrophoresis of PCR products amplified using GFP-specific primers. Lanes 1 and 10: 1 kb^+^ DNA ladder (500 bp band indicated); Lanes 2–3: wild-type samples (no amplification); Lanes 4–7: transgenic lines (distinct ∼500 bp band); Lanes 8–9: PHP81858 plasmid (positive control, 500 bp band), validating primer specificity and PCR conditions. (J) PCR-based validation of Cas9 and g-90RNACas9 integration in transgenic *S. polyrhiza* lines. Agarose gel electrophoresis showing PCR amplification using primers specific to the Cas9 gene (∼700 bp) and guide RNA (gRNA, ∼500 bp). Lanes 1 and 18: 1 kb^+^ DNA ladder with band positions marked for 500 bp and 700 bp. Lanes 2–3: Wild-type (WT) control 1 amplified with Cas9 (Primer 5 and 6 in Table 3) and g-90RNA primers, respectively (Primers 1 and 2 in Table 3); Lanes 4–5: WT control 2 with the same primer sets. No amplification is observed in WT samples, confirming the absence of transgene sequences. Lanes 6–7: g-90RNACas9 transgenic line 1 shows amplification with both Cas9 and gRNA primers, indicating successful integration of both elements. Lanes 8–9: g-90RNACas9 transgenic line 2 shows amplification with Cas9 primers only, suggesting partial or incomplete integration of the gRNA cassette. Lanes 10–11: Cas9 transgenic line 1 exhibits strong amplification with Cas9 primers but no detectable gRNA band. Lanes 12–13: Cas9 transgenic line 2 also shows Cas9 integration without gRNA amplification. Lanes 14–17: Positive control plasmid DNA amplified with Cas9 primers (lanes 14 and 16) shows the expected 700 bp band; no amplification is observed with gRNA primers (lanes 15 and 17), confirming primer specificity and PCR conditions.

**Table 5.**
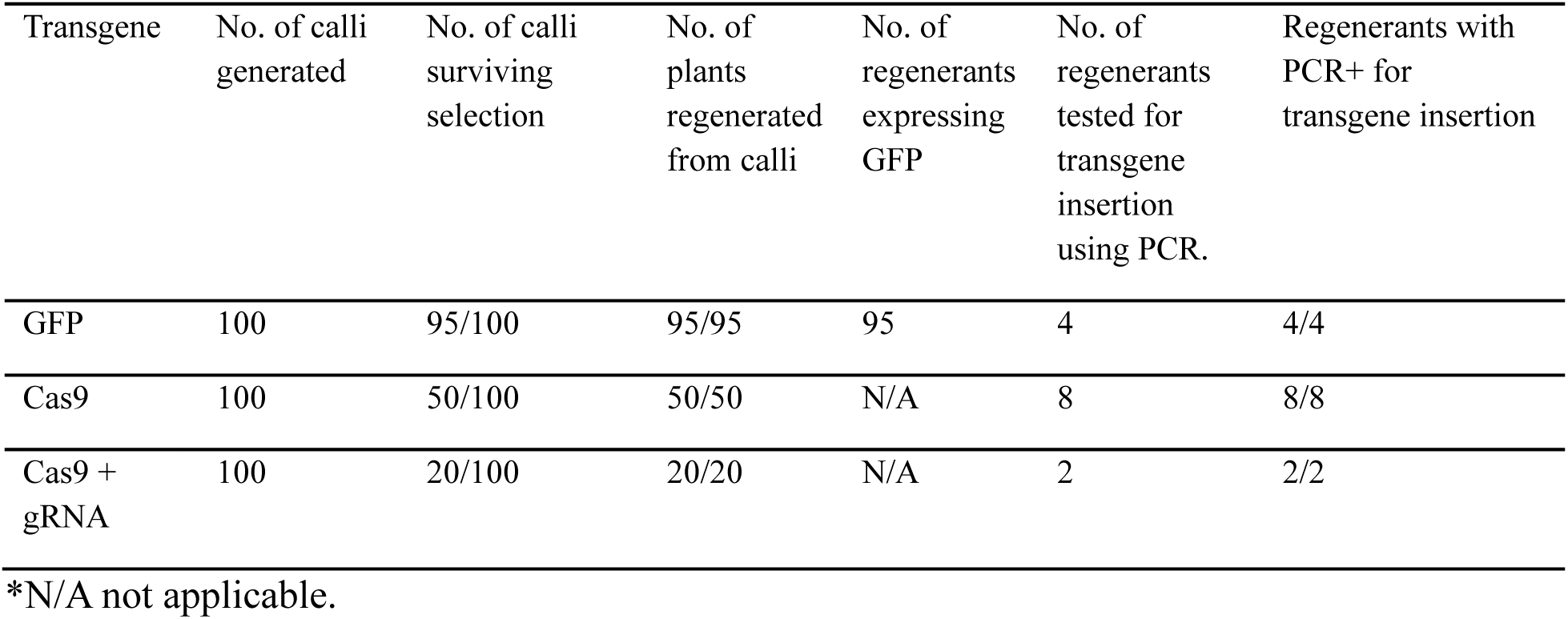
Efficiency of Stable Transformation in *S. polyrhiza*.

| Transgene | No. of calli generated | No. of calli surviving selection | No. of plants regenerated from calli | No. of regenerants expressing GFP | No. of regenerants tested for transgene insertion using PCR. | Regenerants with PCR+ for transgene insertion |
| --- | --- | --- | --- | --- | --- | --- |
| GFP | 100 | 95/100 | 95/95 | 95 | 4 | 4/4 |
| Cas9 | 100 | 50/100 | 50/50 | N/A | 8 | 8/8 |
| Cas9 + gRNA | 100 | 20/100 | 20/20 | N/A | 2 | 2/2 |

\*N/A not applicable.

To mitigate potential phenotypic abnormalities associated with prolonged expression of morphogenic genes, we incorporated an optimized excision step prior to shoot regeneration. The PHP81858 vector includes LoxP sites flanking the *Wus2*, *Bbm*, and *Cre* expression cassettes, allowing their removal prior to T_0_ plant recovery (44). Transformed calli were cultured on ABA (50 µM)-supplemented medium for two days, which induced excision of the morphogenic regulators. Subsequently, G418-resistant calli were transferred to TDZ-containing regeneration medium (R1) and maintained without antibiotics for up to nine weeks (Figure 5F).

Regenerated fronds showed strong and sustained GFP floursecence at 2 weeks (Figures 6C, S8), 6 weeks (Figure 6E), and even 9 weeks (Figures 6G, S9–S11), while no fluorescence signal detected in control plants (Figures S12–S14). Importantly, no abnormal phenotypes were observed in regenerated lines, supporting the effective removal of *Bbm* and *Wus2*.

Notably, GFP cyan fluorescence remained consistently detectable in regenerated *S. polyrhiza* fronds cultured in vitro for more than 270 days (∼115 generations), providing strong evidence for the long-term stability and inheritance of the introduced transgene.

To confirm stable transgene integration, PCR amplification using GFP-specific primers (Primers 9 and 10; Table S2) produced the expected ∼500 bp amplicon in transgenic lines (Figure 6I), but not in wild-type controls. Based on these results, we conclude that the LBA4404 Thy-PHP81858-based transformation system enables efficient, stable, and phenotypically normal transformation of *S. polyrhiza*, achieving a transformation efficiency of 95 transgenic calli per 100 calli (Table 5).

As an additional test of our optimized protocol for plasmids that do not have a visual marker (like GFP), a plasmid containing the gene for CRISPR effector Cas9 was introduced into *S. polyrhiza* calli using *Agrobacterium tumefaciens* strain LBA4404 Thy⁻, harboring the virulence helper plasmid PHP71539 and a Cas9 binary vector which requires hygromycin selection. To our knowledge, no Cas9-expressing strain of *S. polyrhiza* has been generated, which would be useful for high-throughput knockout studies. As before, consistent transformation events were observed when the bacterial suspension was adjusted to an optical density at 600 nm (OD_600_) of 0.8. Following co-cultivation, explants were transferred to selection medium supplemented with 5 mg/L Hygromycin and incubated for three weeks. Hygromycin-resistant calli were subsequently transferred to regeneration medium and cultured for up to nine weeks in the absence of antibiotics (Figure S15), and putative transformants were screened by PCR using Cas9-specific primers to confirm transgene integration (Figure 6J). Of the 100 calli subjected to transformation, 50 survived selection and successfully regenerated, and each of the regenerants tested using PCR were positive for the transgene (Table 5). To evaluate Cas9 nuclease activity, additional transformations were performed using constructs containing guide RNAs targeting the *PDS1* gene (Figure S16); however, no albino phenotypes (Figure S17)—indicative of *PDS1* disruption—were observed no mutations at the target locus were detected among the regenerated lines (S18). While optimization of the CRISPR-Cas9 system in *S. polyrhiza* remains the subject of ongoing optimization (see Discussion below), these results further demonstrate the establishment of a transformation system in *S. polyrhiza* even for plasmids that do not have a visual marker.

## Discussion

Previous studies on *S. polyrhiza* have explored callus induction and demonstrated that the selection and concentration of hormones and carbon sources significantly influence the outcome (30–32). However, despite valuable insights, most of these studies did not fully optimize media compositions for maximum efficiency or take advantage of new advancements such as the use of morphogenic factors to improve stable transformation.

In the current study, we conducted a comprehensive and systematic evaluation of seven culture media (Table 1) to identify optimal conditions for callus induction and regeneration in *S. polyrhiza*. Among these, four media—CIM2, CIM3, CIM5, and CIM6—were previously developed specifically for S. polyrhiza. Reported callus induction efficiencies for these media were 80-100%, not specified, 92%, and 90% respectively. Under our experimental conditions, however, the observed induction efficiencies were 73% for CIM2, 72% for CIM3, 47% for CIM5, and 20% for CIM6, highlighting considerable variability in media performance across experimental setups.

Additionally, we tested two other media originally designed for different species. CIM4, developed for *L. minor*, had no reported induction efficiency, but in our study, it resulted in 63% callus induction. CIM7, developed for *Eragrostis tef*, was previously reported to yield 91.9% efficiency; however, it failed to induce callus formation in *S. polyrhiza* under our conditions.

For regeneration, we tested two media formulations: RM1, previously reported to yield 96% regeneration efficiency, and RM2, reported at 70–80%. In our study, RM1 resulted in 100% regeneration efficiency, while RM2 achieved ≥90%, confirming the high regenerative potential of S. polyrhiza under optimized conditions.

A critical factor influencing callus induction is the developmental stage and morphology of the explant. Our findings confirm that mature fronds with well-developed rhizoids are the most competent for callus formation, consistent with prior observations (45). Notably, 2-week-old explants demonstrated significantly higher callus induction rates across all tested conditions, underscoring the importance of explant maturity and physiological readiness for cellular reprogramming.

The highest callus induction rate (92%) was achieved with CIM1, which contained equal concentrations (0.1 mg/L) of the auxin 2,4-D and the cytokinin TDZ. This balance between auxin and cytokinin proved to be optimal in our system, unlike in other duckweed species such as *S. oligorrhiza*, *S. punctata*, *L. gibba*, *L. minor*, and *W. arrhiza*, where different hormonal ratios were reported to be effective (1,35–37,46–48). High auxin concentrations inhibited frond regeneration, indicating that an excessive auxin-to-cytokinin ratio can suppress vegetative growth, which aligns with earlier findings in duckweed and other monocots (49,50).

Building on the optimized tissue culture system, we established a highly efficient *Agrobacterium*-mediated transformation protocol for *S. polyrhiza*. By employing the LBA4404 Thy⁻ strain harboring the binary vector PHP81858, we achieved a transformation efficiency of 95% at the callus level, with the entire process—from infection to the regeneration of transgenic plants—completed within 5–6 weeks. This represents a significant improvement over previously reported transformation protocols in *S. polyrhiza*, which often required longer durations and yielded lower efficiencies (31,47). Several factors likely contributed to this enhanced efficiency, including the use of a more virulent *Agrobacterium* strain, carefully optimized co-cultivation conditions, and the inclusion of morphogenic regulators *Wuschel2* (Wus2) and *Baby boom* (Bbm) to promote regeneration. The presence of LoxP sites in the vector system allowed for Cre-mediated excision of these regulators prior to regeneration, minimizing the risk of phenotypic abnormalities commonly associated with their constitutive expression. Importantly, we observed sustained GFP fluorescence in regenerated fronds maintained under in vitro conditions for over 270 days (∼115 generations), demonstrating the long-term stability and heritability of transgene expression in *S. polyrhiza* at least in clonally propagating species.

The establishment of an efficient transformation system enabled us to perform CRISPR/Cas9-mediated genome editing in *S. polyrhiza*. As a proof-of-concept, we targeted the phytoene desaturase (*SpPDS*) gene, a commonly used visual marker for genome editing due to the albino phenotype resulting from its disruption(51). Despite successful integration of Cas9 and gRNA constructs (as confirmed by PCR), none of the regenerated plants displayed the expected mutant phenotype or sequence-level edits. There are several plausible explanations for the lack of detectable editing. First, the gRNA may have exhibited low cleavage efficiency due to suboptimal target sequence characteristics, such as secondary structure or low chromatin accessibility. Second, the *SpPDS* locus may be embedded in tightly packed heterochromatin, which can inhibit the access and activity of the CRISPR/Cas9 complex. Chromatin structure is known to influence gene editing efficiency, especially in species with compact genomes such as *S. polyrhiza*. Finally, mutations may have occurred at low frequencies or in chimeric tissues, making them undetectable by conventional PCR and Sanger sequencing. Future work should consider using multiple gRNAs, protoplast-based assays, and high-throughput sequencing to improve mutation detection and confirm editing efficiency.

We also successfully established a highly efficient transient transformation protocol in *S. polyrhiza* by optimizing vacuum infiltration, a post-infiltration resting period, and acetosyringone supplementation during both inoculation and co-cultivation. These parameters significantly enhanced gene delivery and expression, as confirmed by robust GUS reporter activity in fronds transformed using *Agrobacterium* strain GV3101. In contrast, strain LBA4404 failed to mediate successful transformation, underscoring the critical role of strain selection in determining transformation efficiency. This finding is consistent with previous reports in other plant systems, where differences in virulence genes, host range, and T-DNA delivery efficiency between *Agrobacterium* strains have been shown to influence transformation outcomes (52,53). For example, GV3101 has been successfully used in various monocots and dicots due to its higher virulence and compatibility with a broad range of plant tissues (54). Although transient transformation in duckweed has been demonstrated previously (33), our study presents an improved and reproducible protocol specifically tailored for *S. polyrhiza*, with transformation efficiency exceeding 90%. These results provide a valuable framework for rapid gene expression studies in duckweed and offer a basis for downstream applications such as CRISPR/Cas9-based functional genomics. In fact, we expect that combination of the approaches presented here, along with our previously-reported carbon nanotube (CNT)-mediated plasmid DNA delivery system (9) could be used for rapid, transient delivery of plasmid DNA such that by combining transient and stable delivery modes, short-term screening of editing efficiency and the generation of stable transformants carrying validated CRISPR components can be readily achieved.

In conclusion, we present a streamlined pipeline for tissue culture, genetic transformation, and genome editing in *S. polyrhiza*, a representative of an early-diverging monocot lineage. With a small nuclear genome (∼158 Mb) and the most compact mitochondrial genome reported among flowering plants, *S. polyrhiza* offers a powerful system for molecular and synthetic biology. The successful development of callus induction, regeneration, and genetic transformation protocols not only advances the functional study of duckweed biology but also accelerates its application in bioenergy, wastewater remediation, and plant-environment interaction studies as well as high-throughput platform for its myriad emerging applications in functional genomics, synthetic biology, and plant biotechnology

## Materials and Methods

### Plant Materials and Cultivation Conditions

*Spirodela polyrhiza* 9509 was obtained from the Rutgers Duckweed Stock Cooperative (ruduckweed.org). Unless otherwise specified, *S. polyrhiza* was used as the primary experimental material. The duckweed was collected from agar plates, surface sterilized with 0.5% sodium hypochlorite and rinsed thoroughly with autoclaved water. Following sterilization, the plants were cultivated in a cost-effective hydroponic system, where they were grown in a nutrient-rich liquid medium instead of agar. The system consisted of two 3-liter stackable bins placed in parallel across a 14-liter plastic reservoir tub. The bins were secured using their lips, leaving a 6-inch void space in the reservoir below. Each bin was modified with a nutrient feed pipe at the bottom, allowing drainage into the reservoir. A standard 3W, 50 gph submersible aquarium pump was placed in the reservoir and fitted with a Y-shaped tube to circulate water between the reservoir and the stacked bins. To prevent algae growth, a UV sterilization unit (Green Killing Machine, Internal UV) was installed in the reservoir. The hydroponic system was filled with commercially available nutrient solutions (General Hydroponics FloraSeries, including FloraMicro, FloraBloom, and FloraGro) at a 1:1 ratio. Water circulation and aeration were maintained continuously. To prevent overcrowding, plant density was adjusted weekly, and nutrient renewal was determined by monitoring conductivity. The plants were illuminated using a full-spectrum LED grow light positioned 200 mm above the growth area, following a 16-hour light/8-hour dark cycle. The system was maintained under controlled environmental conditions, including nutrient availability, temperature, light intensity, and photoperiod, to optimize growth.

*S. polyrhiza* was prepared for the experiment by undergoing surface sterilization with 2.5% sodium hypochlorite, treating *S. polyrhiza* for 3 minutes followed by 5–8 rinses with autoclaved water. The plants were then cultured in Magenta jars containing 50 mL of half-strength Schenk & Hildebrandt (0.5× SH) basal salt medium, supplemented with 1% (m/v) sucrose at pH 5.8. Cultures were maintained in plant growth chamber at 28°C under an irradiance of 85 μmol photons PAR m⁻² s⁻¹, with a 16-hour light/8-hour dark cycle, and subcultured weekly. To determine the optimal conditions for callus induction, fronds were cultivated on seven different callus induction media (CIM) (Table 1). Mature fronds with well-developed rhizoids were selected from aseptic cultures, chopped, and transferred into petri dishes. Approximately 20 explants were placed in sterilized 9 cm petri dishes containing 40 mL of solid callus induction medium. These explants were incubated in the dark at 28°C in an artificial plant growth chamber for five weeks. The number of explants that successfully induced callus formation was recorded.

### Tissue culture and regeneration

#### Callus Line Establishment and Sustained Growth Maintenance

For callus induction, explants were prepared from sterile in vitro fronds transferred to CIM1, and maintained under dark conditions in the plant growth chamber. Subculturing was performed every two weeks, with growth progress observed and recorded weekly.

#### Plant Regeneration

*S. polyrhiza* callus was transferred to regeneration media (RM) (RM1(31): 0.5X MS medium supplemented with 1% (m/v) sucrose, 0.6% (m/v) agar, 4.54 μMTDZ, pH 5.8 and RM2 (32): SH media, 10 μM zeatin, 1% sucrose, 0.6% (m/v) agar, pH 5.8) cultured at 28 °C in plant growth chamber under an irradiance of 85 μmol photons PAR m⁻² s⁻¹, with a 16-hour light/8-hour dark cycle. These calli with or without regenerated fronds were subcultured every 2 weeks. The state of regeneration was observed and recorded every week. Clustered fronds were transferred to liquid 0.5X SH basal salt media, supplemented with 1% (m/v) sucrose at pH 5.8 containing 1% sucrose until rhizoids initiation.

### Transient transformation

#### Agrobacterium-Mediated Transient Transformation of *S. polyrhiza*

Transient transformation of *S. polyrhiza* was conducted according to Peterson et al. 2021 with modification (33). The binary vector pSB161 (pL2_pSB90_2×35S::GUS::tMAS), which harbors a GUS reporter gene with introns under the control of a double 35S promoter, was obtained from Erin Cram and Carolyn Lee-Parsons (Addgene plasmid #123197; http://n2t.net/addgene:123197; RRID:Addgene_123197). *Agrobacterium* strains GV3101 and LBA4404 (GoldBio) were transformed with pSB161 via electroporation following the manufacturer’s protocol. Transformed *Agrobacteria* were cultured in 150 mL of LB medium supplemented with gentamycin (30 mg/L), rifampicin (25 mg/L), kanamycin (50 mg/mL), and 200 µM AS at 28 °C in 500 mL flasks shaking at ∼180 rpm for 48 h. Bacterial cells were harvested by centrifugation at 4,000 g for 10 min at room temperature and resuspended in an equal volume of 10 mM Mg-MES buffer (pH 5.6) containing 200 µM AS. To ensure submersion of duckweed fronds—which typically float—plants were enclosed in autoclaved, reticulated stainless-steel containers (tea infusers) and immersed in the *Agrobacterium* suspension. Approximately 50 fronds were infiltrated under vacuum (pressure below –0.1 MPa) for 30 min, followed by a 30 min rest at atmospheric pressure. After infiltration, fronds were briefly rinsed with autoclaved water, blotted dry on sterile filter paper in Petri dishes, and incubated in the dark at 25 °C for 24 h in the presence of 200 µM AS. Following co-cultivation, fronds were transferred to SH liquid and solid (0.6% agar) medium supplemented with 30 g/L sucrose and 600 mg/L timentin to eliminate residual *Agrobacterium*. Plants were then incubated under a 16 h/8 h light/dark photoperiod at 25 °C (light) and 16 °C (dark) in a plant growth chamber for further development.

#### GUS Histochemical Assay

*GUS* activity in transiently transformed *S. polyrhiza* fronds was assessed following the protocol described by Yang et al. 2018.(31) Fronds were vacuum infiltrated for 1 h in *GUS* staining solution containing 100 mM sodium phosphate buffer (pH 7.0), 0.5 M EDTA, 50 mM potassium ferricyanide [K_3_Fe(CN)_6_], 50 mM potassium ferrocyanide [K_4_Fe(CN)_6_], 0.1% Triton X-100, and 0.5 mg/mL X-Gluc (5-bromo-4-chloro-3-indolyl β-D-glucuronide), and incubated overnight at 37 °C in the dark. The next day, stained fronds were rinsed with 100 mM phosphate buffer and deionized water. To remove chlorophyll and enhance stain contrast, samples were sequentially treated at 57 °C for 15 min each in: (1) 0.24 N HCl with 20% (v/v) methanol, and (2) 7% (w/v) NaOH with 10% (v/v) ethanol. Final rinses were performed with 40% ethanol, and samples were stored in a solution of 5% (v/v) ethanol and 25% (v/v) glycerol. GUS activity was visualized using a light microscope. The presence of dark blue spots indicated successful transient expression of the GUS reporter. Fronds exhibiting one or more blue foci were scored as “*GUS* positive.”

### Stable transformation

#### Preparation of CRISPR gRNA construct

*PDS* gene, which encodes phytoene desaturase—a widely used marker for assessing gene-editing efficiency in plants (55,56). The *PDS* gene from *Oryza sativa* was used as a reference and subjected to CoGe BLAST against the *S. polyrhiza* genome. A homologous coding region was identified on Chromosome 18, spanning positions 60,549 to 68,089. This sequence was analyzed using the CRISPOR online tool, which identified four best candidate guide RNAs (gRNA) target sites within the *SpPDS* genomic region (Table S1). Each gRNA was individually cloned into the binary Cas9 expression vector JD633 following the Joung Lab protocol (Addgene guide; https://media.addgene.org/data/plasmids/65/65768/65768-attachment_B-IZ6VSu18MX.pdf). The JD633 vector backbone was digested using the AarI restriction enzyme (https://www.thermofisher.com/order/catalog/product/ER1581). Successful integration of gRNA sequences was confirmed by PCR analysis followed by sanger sequencing (Figure S16).

#### *Agrobacterium*-Mediated Stable Transformation of *S. polyrhiza*

For *Agrobacterium*-mediated stable transformation of *S. polyrhiza*, we followed a modified version of the protocol by Yang et al. 2018 (31). The transformation was performed using *Agrobacterium* strain LBA4404 Thy⁻, containing the vir helper plasmid PHP71539 obtained from Corteva Agriscience. This host strain was transformed with the binary vectors PHP81858, also obtained from Corteva Agriscience and JD633 (with or without guide RNAs), the latter binary vector is available at Addgene (#160393). Transformation of the host strain was carried out using a heat shock protocol (GoldBio protocol).

#### Preparation of *Agrobacterium*

*Agrobacterium* (LBA4404 Thy⁻) strains harboring the desired plasmids were cultured on solid Yeast Extract Broth (YEB) medium (0.6% yeast extract, 0.5% tryptone, 0.5% glucose, 0.6% agar, and 2 mM MgSO_4_; pH 7.0), supplemented with the appropriate antibiotics. For JD633, the medium was supplemented with thymidine (50 mg/L), gentamycin (50 mg/L), and kanamycin (50 mg/L). For PHP81858, thymidine (50 mg/L), gentamycin (50 mg/L), and spectinomycin (50 mg/L) were added. Cultures were incubated at 28 °C for 2 days. Colonies were then inoculated into liquid YEB medium with the same antibiotics and grown until the optical density at 600 nm (OD_600_) reached 0.8.

### Co-cultivation

Light yellow calli were sectioned into 2–3 mm pieces and immersed in the *Agrobacterium* suspension. The samples were shaken at 100 rpm for 20 minutes, followed by vacuum infiltration at 0.8 kg/cm² for 10 minutes. Treated calli were transferred to co-cultivation medium (0.5X MS, 4.58 mM MES, 22.62 μM 2,4-D, 8.88 μM 6-BA, 100 μM AS, and 50 mg/L thymidine, pH 5.2).

Co-cultivation was performed in 9 cm Petri dishes containing six layers of sterilized filter paper soaked in 6 mL of the liquid medium. Plates were incubated in the dark at 25 °C for 5 days, with 8 calli per plate. All experiments were conducted in triplicate.

### Resting and Selection

Following co-cultivation, calli were transferred to resting medium (0.5X MS with vitamins, 1% sucrose, 0.6% agar, 9.99 mM (NH_4_)_2_SO_4_, 22.62 μM 2,4-D, 8.88 μM 6-BA, and 0.63 mM cefotaxime; pH 5.8) and incubated for 4 days. Calli were then transferred to selection media depending on the plasmid. For PHP81858: 0.5× MS medium containing 1% sucrose, 0.6% agar, 9.99 mM (NH_4_)_2_SO_4_, 22.62 μM 2,4-D, 8.88 μM 6-BA, 0.63 mM cefotaxime, and 30 mg/L G418 (pH 5.8). For JD633: 0.5× MS medium with the same components but supplemented with 5 mg/L hygromycin instead of G418. Selection was carried out over three weeks, with weekly subculturing on fresh medium. Regeneration and selection were conducted at 28 °C in the dark.

### Estimation of Transformation Efficiency (TE) and Visualization of Reporter genes

After 1 month of selection, all resistant calli expressing the AmCyan1 from each petri dish were considered as a single transgenic event. The transformation efficiency (TE) was calculated as percentages of resistant calli in the total number of explants. The Cyan florescence was visualized using Nikon SMZ18 fluorescent microscope using a GFP filter.

### Excision of *Bbm* and *Wus2* Genes

Because the morphogenic genes Bbm and Wus2 may produce undesirable genotypes, calli transformed with the PHP81858 binary vector were transferred to excision medium (0.5X MS with 1% sucrose, 0.6% agar, 9.99 mM (NH_4_)_2_SO_4_, 22.62 μM 2,4-D, 8.88 μM 6-BA, and 50 μM ABA; pH 5.8) and incubated at 28 °C in the dark to induce excision of the *Wuschel2* (*Wus2*) and *Baby Boom* (*Bbm*) genes.

### Shoot Induction and Plant Regeneration

Antibiotic resistant transgenic calli were transferred to shoot induction medium (0.5X MS supplemented with 1% sucrose, 0.6% agar, and 4.54 μM TDZ; pH 5.8), with weekly subculturing. Regenerated fronds from different transformation events were isolated and cultured individually in liquid 0.5X MS medium (pH 5.8) containing 1% sucrose. Cultures were maintained at 28 °C under 85 μmol photons m⁻² s⁻¹ PAR with a 16 h photoperiod.

### PCR Analysis

To confirm the presence of guide RNAs in the JD633 plasmid, bacterial colony PCR was performed. Primers 1 and 2 (Table 3) were used for this purpose. Each 25 μL reaction contained 12.5 μL of OneTaq® 2X Master Mix with Standard Buffer (New England Biolabs), 1.0 μL each of 10 μM forward and reverse primers, 11.5 μL deionized water, and a single bacterial colony as the DNA template. Amplification was carried out in a thermal cycler (Eppendorf, USA) using the following program: initial denaturation at 94 °C for 30 s; 35 cycles of denaturation at 94 °C for 30 s, annealing at 50 °C for 60 s, and extension at 68 °C for 1 min; followed by a final extension at 68 °C for 5 min. Amplified products were separated on a 1% (w/v) agarose gel and visualized using a UV transilluminator. Selected amplicons were purified and submitted for sequencing (Azenta Life Sciences, USA) to confirm accurate guide RNA insertion.

To verify the successful transformation of *Agrobacterium* strains with the binary plasmids, colony PCR was performed using primers 3 and 4 for the PHP81858 plasmid. For the JD633 plasmid, primers 5 and 6 and primers 7 and 8 were used (Table 3).

For transgenic plant analysis, genomic DNA was extracted from regenerated plants using the Wizard® HMW DNA Extraction Kit (Promega, USA), following the manufacturer’s protocol. Genomic DNA (50 ng per reaction) was used as the template in a 25 μL PCR reaction composed of 12.5 μL GoTaq® Master Mix (Promega), 1.0 μL each of 10 μM forward and reverse primers, and 10.5 μL of deionized water. PCR confirmation of guide RNAs in transgenic plants used the same primers as the bacterial screening. To verify the presence of the PHP81858 T-DNA construct in transgenic lines, the primers 9 and 10 were used (Table 3). Amplification was performed in a thermal cycler (Applied Biosystems, USA) with the following cycling conditions: initial denaturation at 95 °C for 2 mins; 30 cycles of 95 °C for 1 min, 55 °C for 45 s, and 72 °C for 45 s; followed by a final extension at 72 °C for 5 min. PCR products were resolved on 1% (w/v) agarose gels and visualized using UV transillumination.

## Statistical analysis

The callus induction and regeneration experiments were conducted with a minimum of five replicates, each consisting of 20 explants and 10 calli. All experiments were repeated three times. Results are expressed as means ± standard deviations. The efficiency of callus induction on each plate was assessed after one month of cultivation, while the frond regeneration rate was evaluated after five weeks. Statistical significance was determined using unpaired t-tests.

## Supplementary information

A supplementary document associated with this manuscript contains additional images of duckweeds references in the main text.

## Abbreviations

MS: Murashige and Skoog
SH: Schenk and Hildebrandt
2,4-D: 2,4 Dichlorophenoxyacetic acid
TDZ: Thidiazuron
6-BA: 6-Benzylaminopurine
ABA: Abscisic acid
NAA: 1-Naphthaleneacetic acid
AS: Acetosyringone
MES: 2-(N-morpholino) ethanesulfonic acid

## Acknowledgments

We thank members of the Josephs laboratory, particularly Dr. Rachel Tinker-Kulberg for helpful discussions regarding duckweed maintenance. The work was funded by grants from the National Institute of General Medical Sciences [R35GM133483 to EAJ]. The content is solely the responsibility of the authors and does not necessarily represent the official views of the National Institutes of Health. The work was also supported by funds from the SUNY Empire Innovation Program (EIP) and departmental funds from the Department of Biomedical Engineering at Stony Brook University. The work was also supported in part by the ICONS program at JSNN [US Department of Defense Contract #W911QY2220006]. The work was also supported by the NSF Plant Genome Research Program (PGRP) [Award ID 2327906 to ALO]. We also acknowledge Corteva Agriscience for their donation of *Agrobacterium* strains and plasmids. This work in part was performed at JSNN, a member of the Southeastern Nanotechnology Infrastructure Corridor (SENIC) and National Nanotechnology Coordinated Infrastructure (NNCI), which is supported by the National Science Foundation [Grant ECCS-1542174].

## Declaration of Interests

Authors have no interests to declare. Corteva Agriscience provided *Agrobacterium* strain LBA4404-Thy-with helper plasmid and the binary vector PHP81858.

