## Supplementary Information for "Systematic Optimization Enables Near-Perfect In Vitro Transformation Efficiencies for *Spirodela polyrhiza* (Greater Duckweed)"

**Figure S1. Additional Images of Calli of *S. polyrhiza* on CIM1**

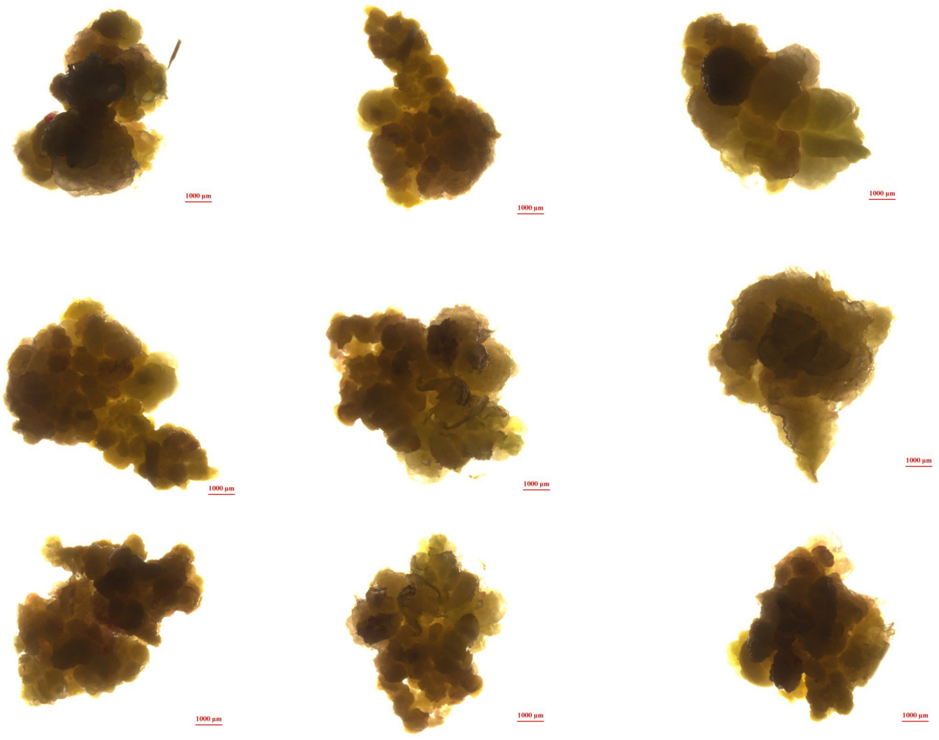

**Figure S2. Additional Images of Proliferating Calli of *S. polyrhiza* on CIM1**

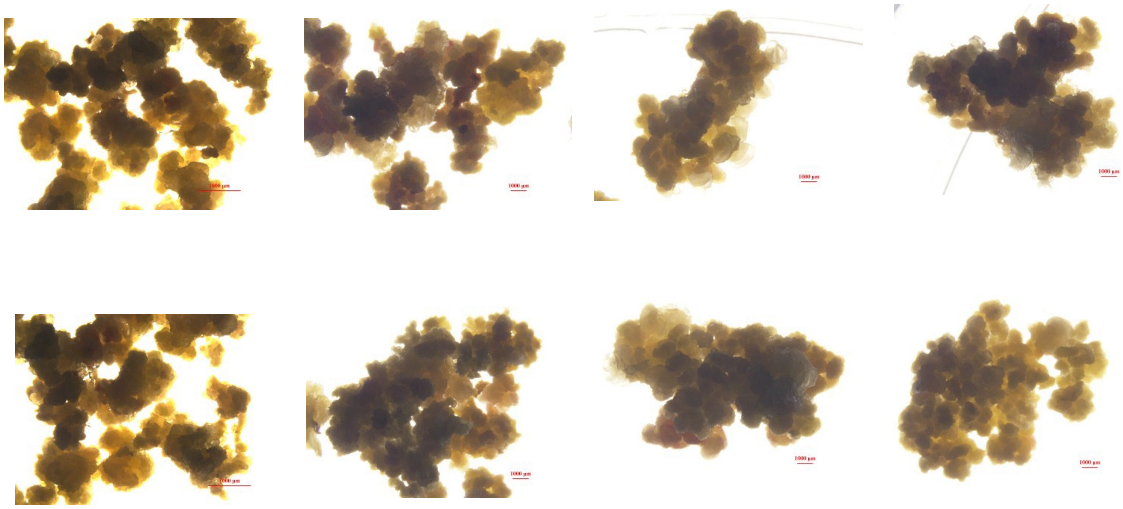

**Figure S3. Additional Images of Regenerated Calli of *S. polyrhiza* on Regeneration Media 1 (R1)**

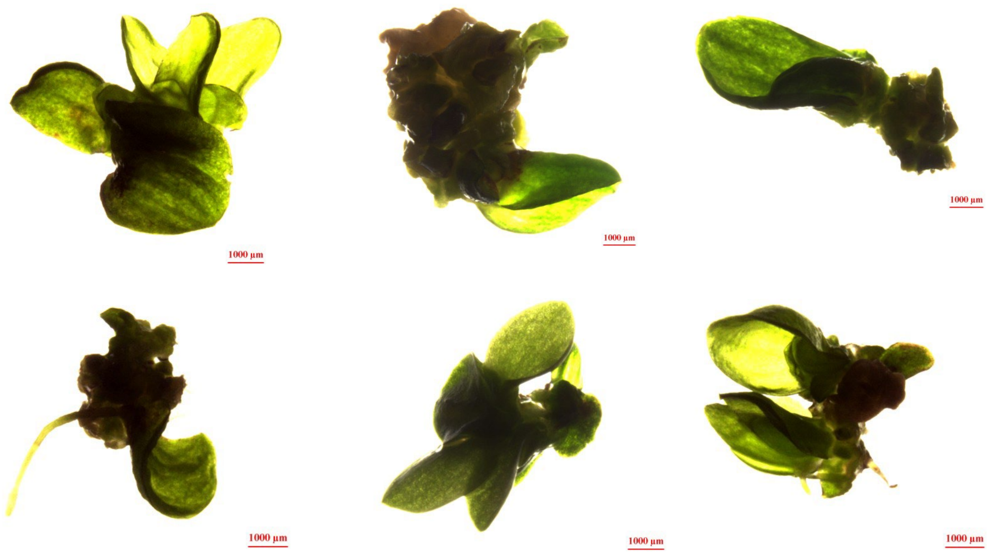

**Figure S4. Additional Images of GUS Positive of *S. polyrhiza* Fronds after GUS Assay**

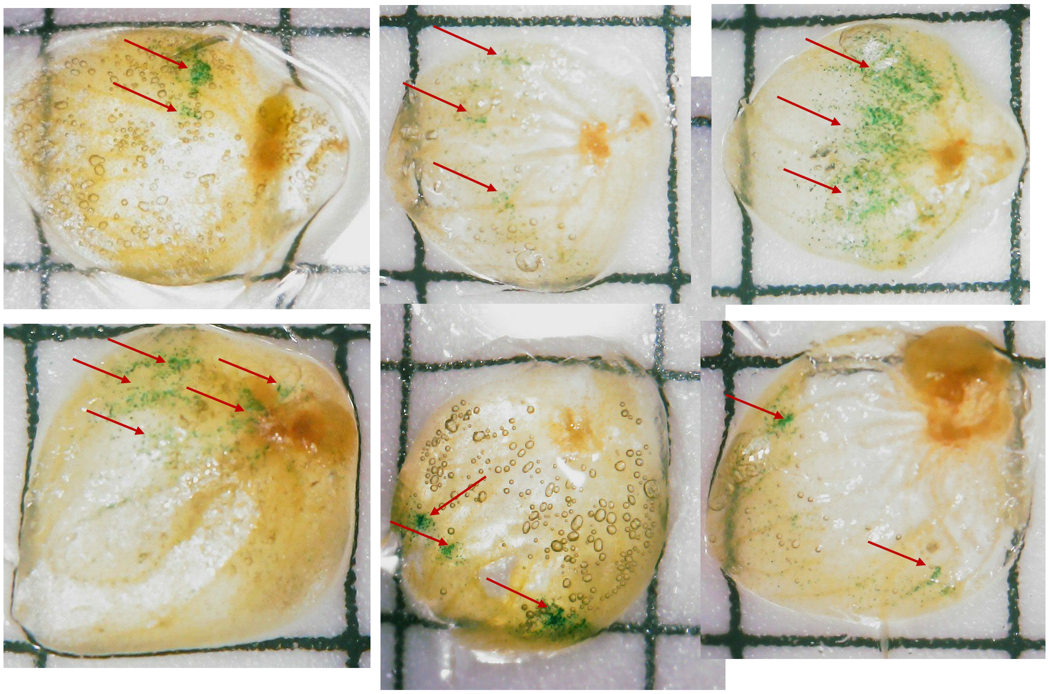

Note. Red arrows in the picture indicate the GUS expression. The grid size in the images is 5 mm.

**Figure S5. Additional Images of Control *S. polyrhiza* Fronds after GUS Assay**

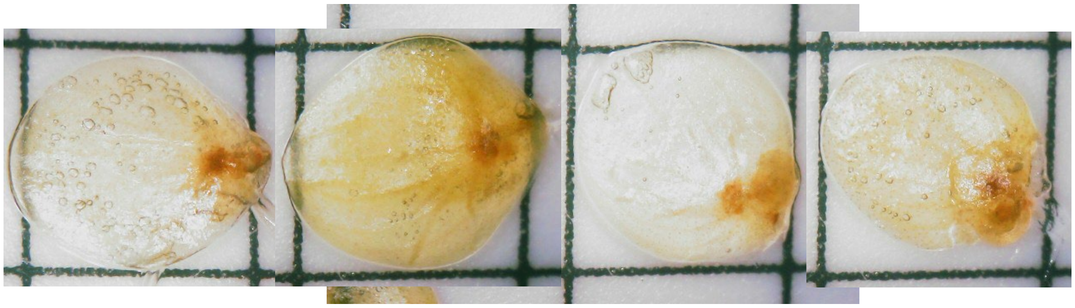

*Note*. Duckweeds were transformed with only GV3101. After GUS staining, none exhibit the characteristic blue regions expected for a positive GUS stain as in Figure 4B.

**Figure S6. Transformed Calli of *S. polyrhiza* Showing GFP Expression on Selection Media in First Week on Selection Media**

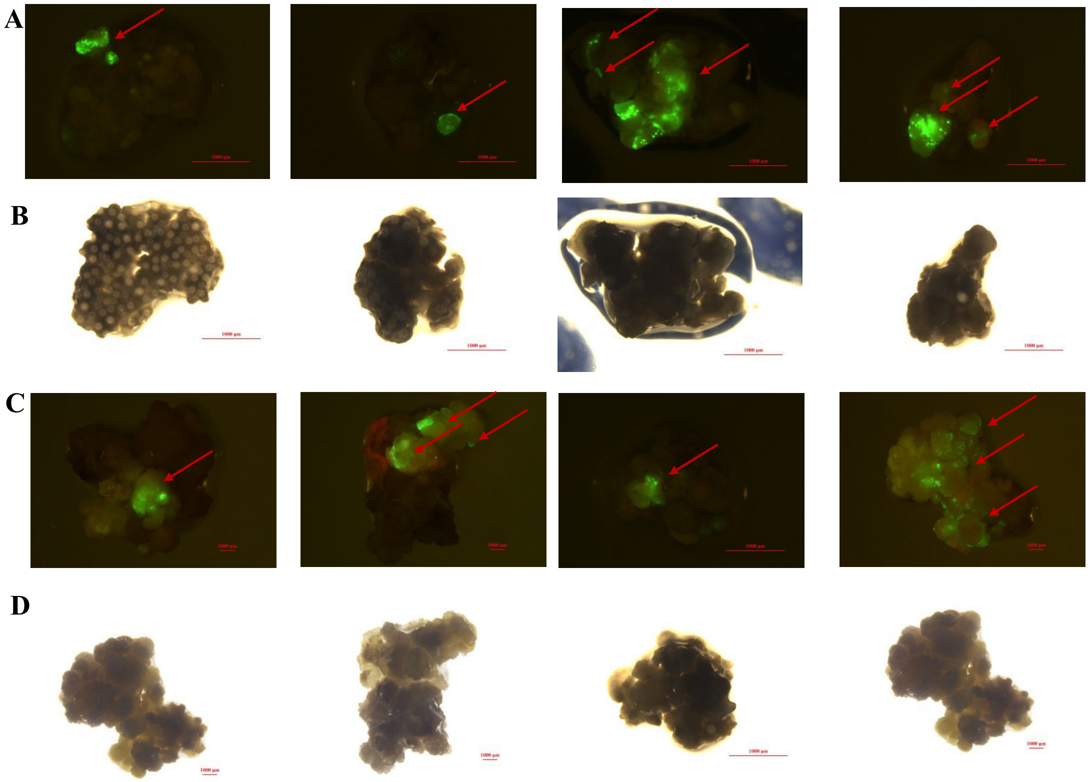


Note. Panels (A) and (C) were captured under blue light to visualize GFP fluorescence, while panels (B) and (D) show the corresponding bright-field images. Red arrows indicate regions of GFP expression.

**Figure S7. Additional Images of Control Calli of *S. polyrhiza* Showing No GFP Expression on Selection Media**

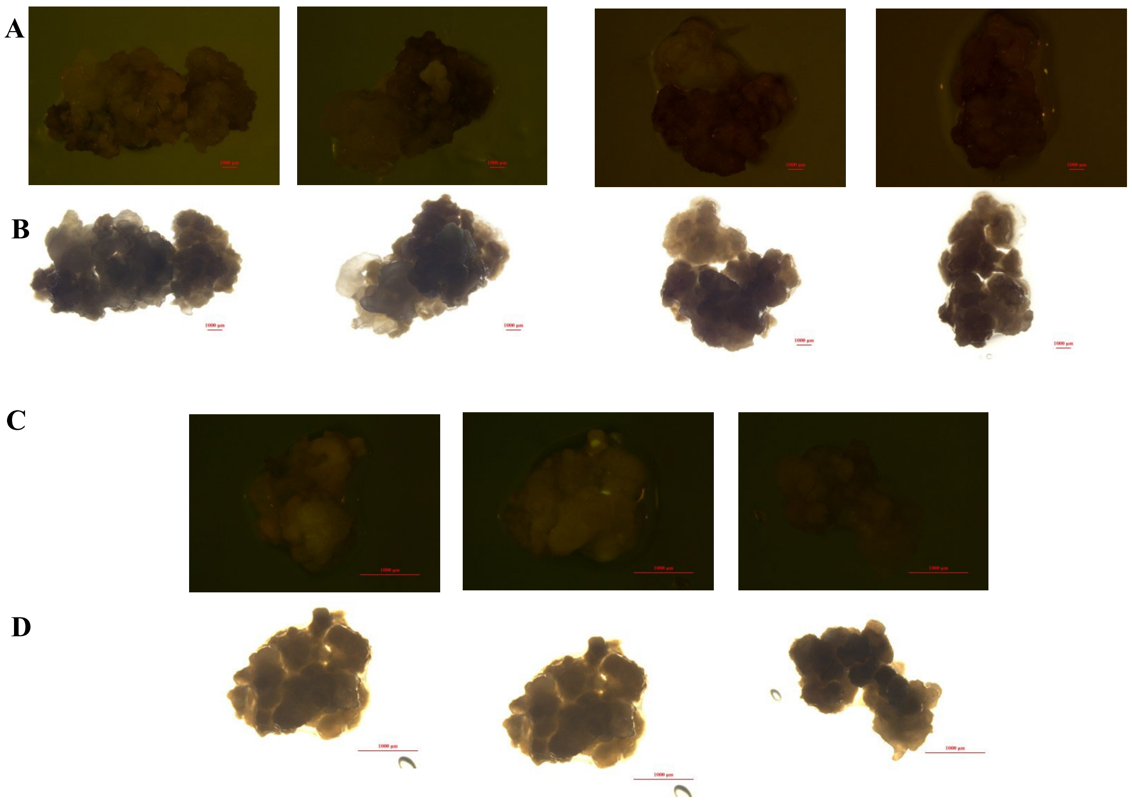


*Note.* Panels (A) and (C) were captured under blue light to visualize GFP fluorescence, while panels (B) and (D) show the corresponding bright-field images. No GFP expression was observed in the control samples.

**Figure S8. Transformed Regenerated calli of *S. polyrhiza* Showing Bright GFP Expression on Regeneration Media After Two Weeks**

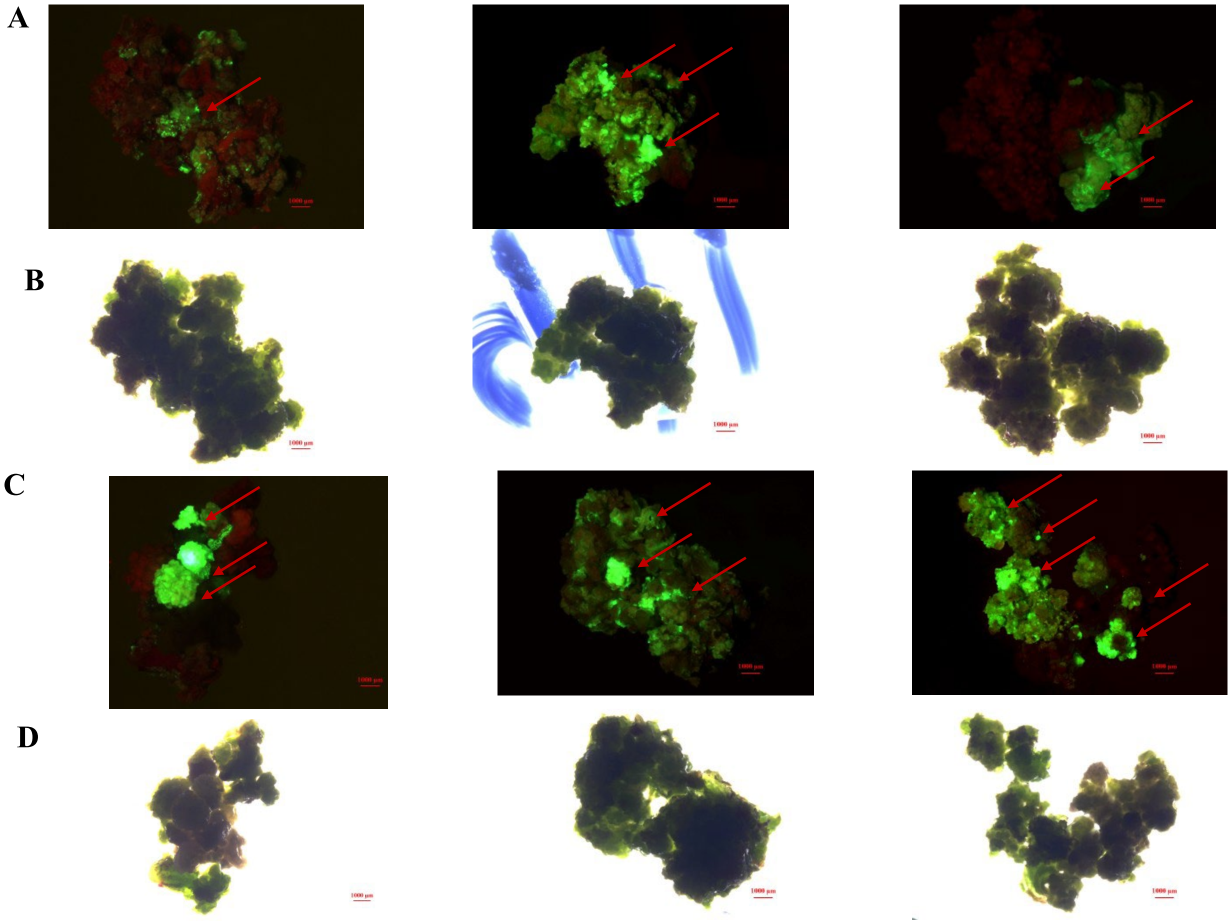

*Note.* Panels (A) and (C) were captured under blue light to visualize GFP fluorescence, while panels (B) and (D) show the corresponding bright-field images. Red arrows indicate regions of GFP expression

**Figure S9. GFP-Expressing Transgenic *S. polyrhiza* Fronds Regenerating on Regeneration Media after Nine Weeks**

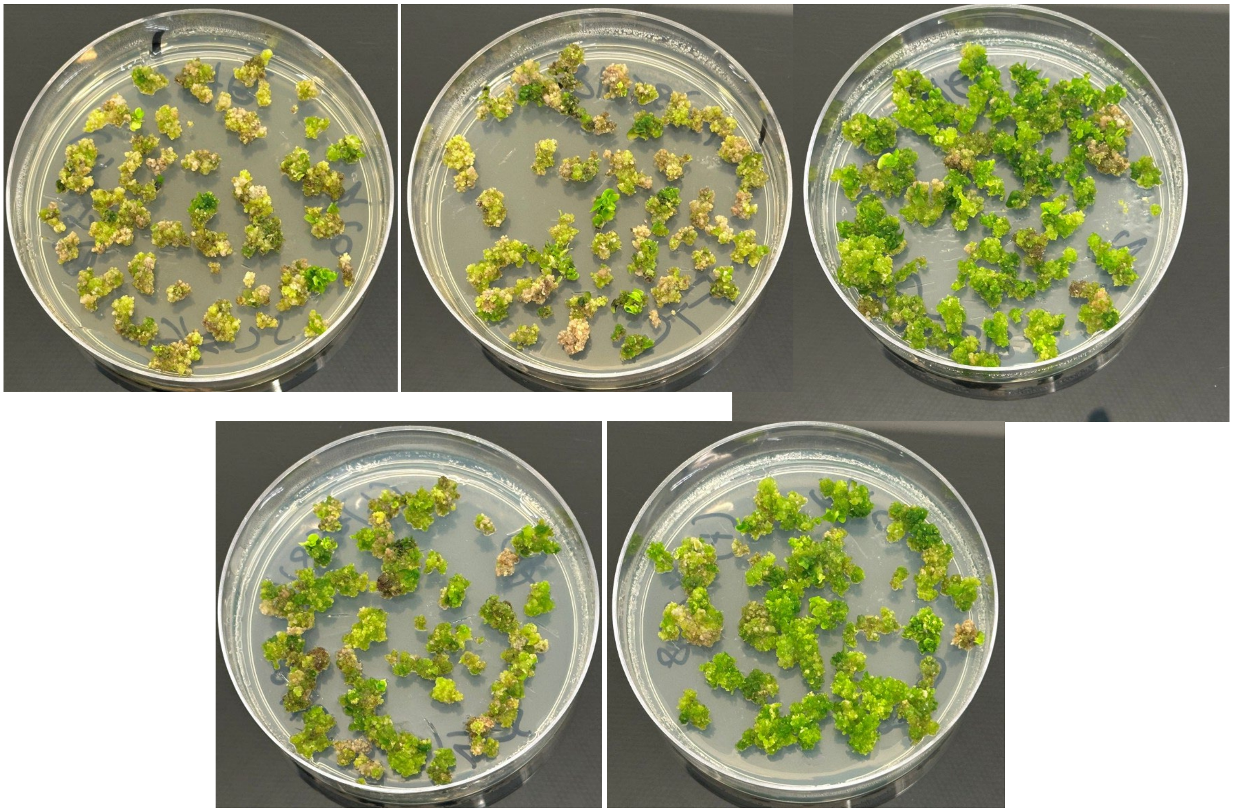

*Note.* Calli were cultured on regeneration medium following selection and showed progressive greening and shoot formation over time. Visible differences in the extent of regeneration and chlorophyll accumulation are observed across different plates, reflecting the high efficiency and consistency of the regeneration process. The green patches indicate emerging shoot structures and successful regeneration. All plates represent independent biological replicates maintained under identical conditions (28 °C, 16/8 h light/dark photoperiod). Scale bars are not shown; petri dish diameter is 90 mm.

**Figure S10. Microscopic Images of *S. polyrhiza* Transgenic GFP Regenerants after Nine Weeks**

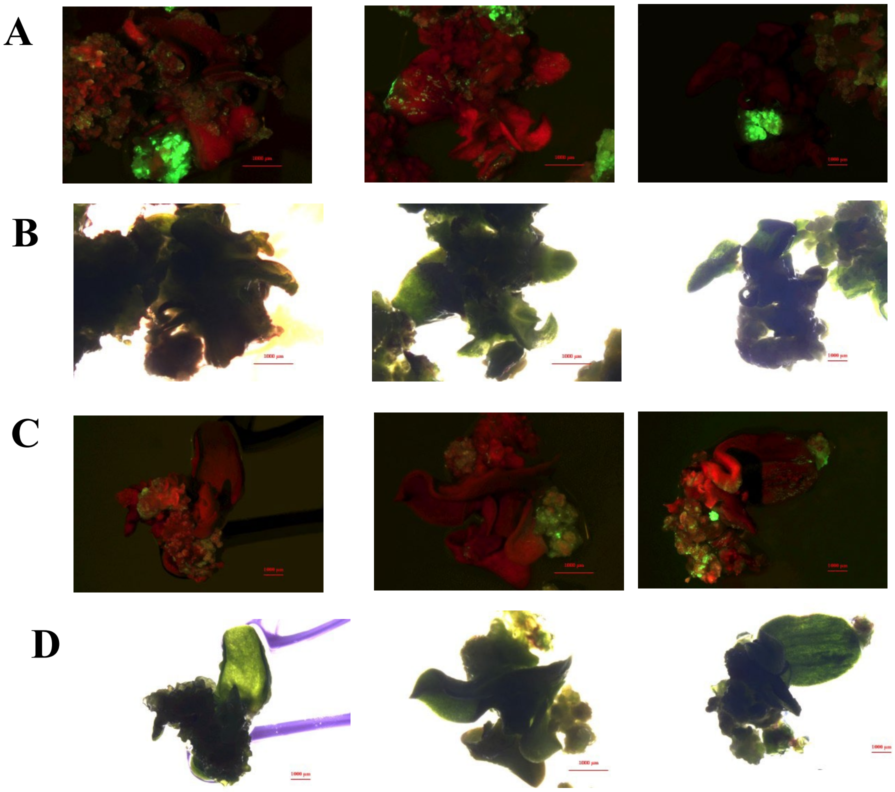


*Note*. (A, C) Bright GFP expression observed in the Transgenic *S. polyrhiza*. (B, D) Corresponding bright-field images of the same GFP-expressing fronds shown in (A, C).

**Figure S11. Additional Images of Regenerating Transgenic GFP plants on Solid Regeneration Media**

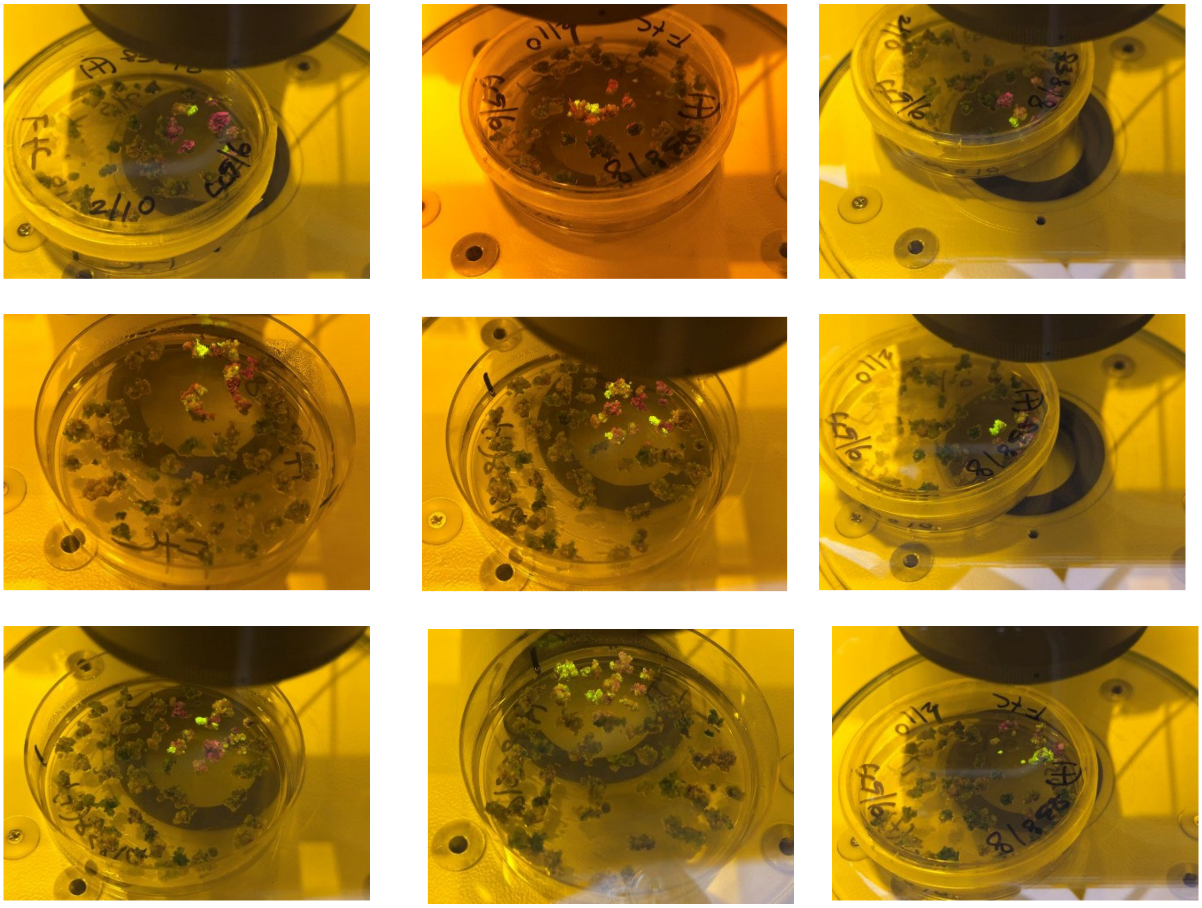

**Figure S12. Additional Images of Control Calli of *S. polyrhiza* Showing No GFP Expression on Regeneration Media**

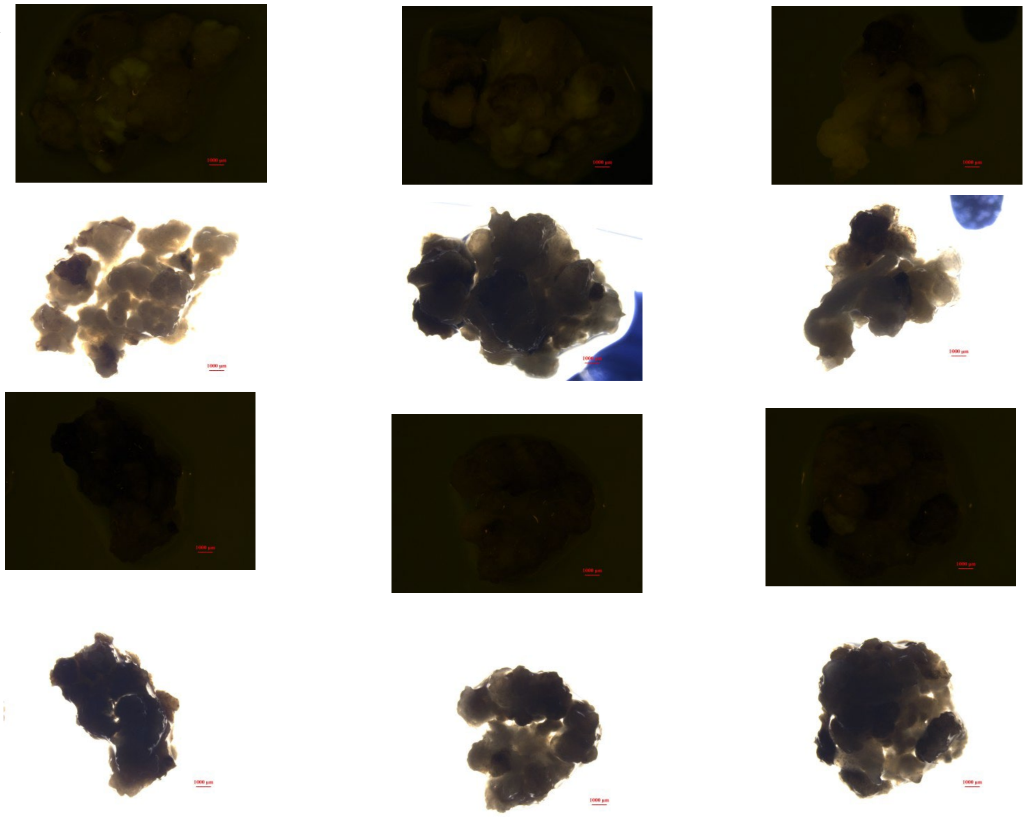

*Note*. Panels (A) and (C) were captured under blue light to visualize GFP fluorescence, while panels (B) and (D) show the corresponding bright-field images. No GFP expression was observed in the control samples.

**Figure S13. Additional Images of Regenerating Wild-Type Control Plants on Solid Regeneration Media**

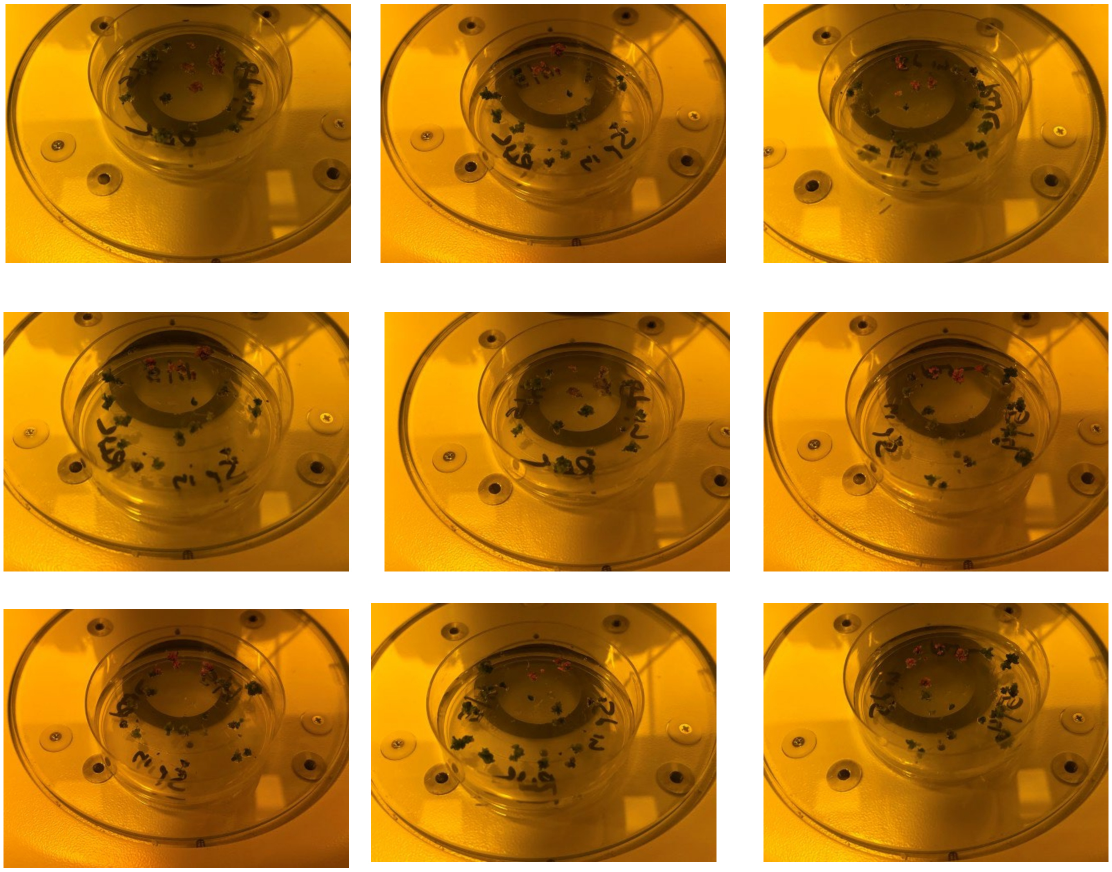

**Figure S14. Additional Microscopic Images of Wild Type *S. polyrhiza* Plant**

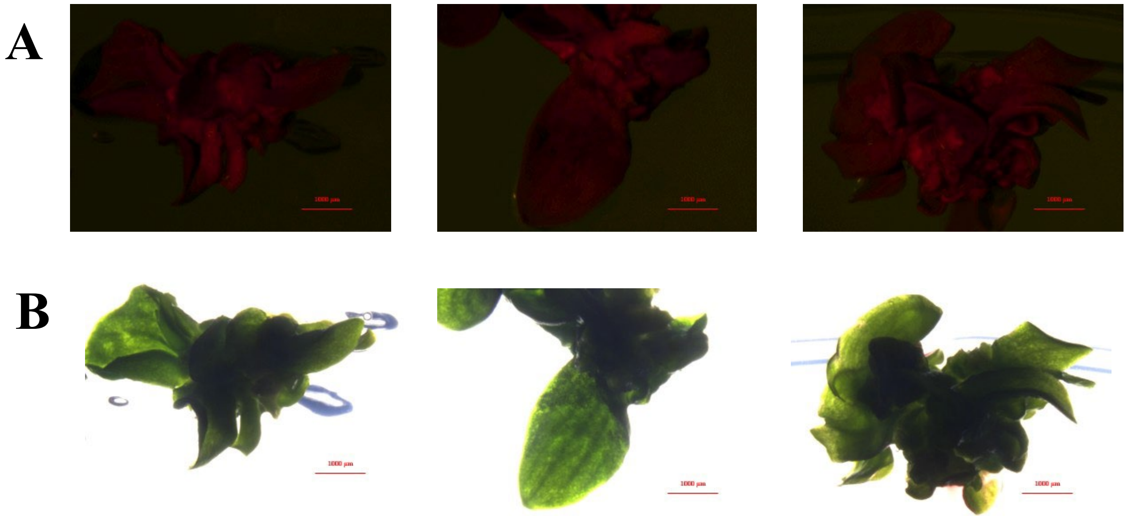

*Note.* (A) No GFP expression observed in the wild type of *S. polyrhiza.* (B) Corresponding bright-field images of the same plants shown in (A).

**Figure S15. Additional Images of Regenerated Fronds of *S. polyrhiza* Containing Cas9 Construct**

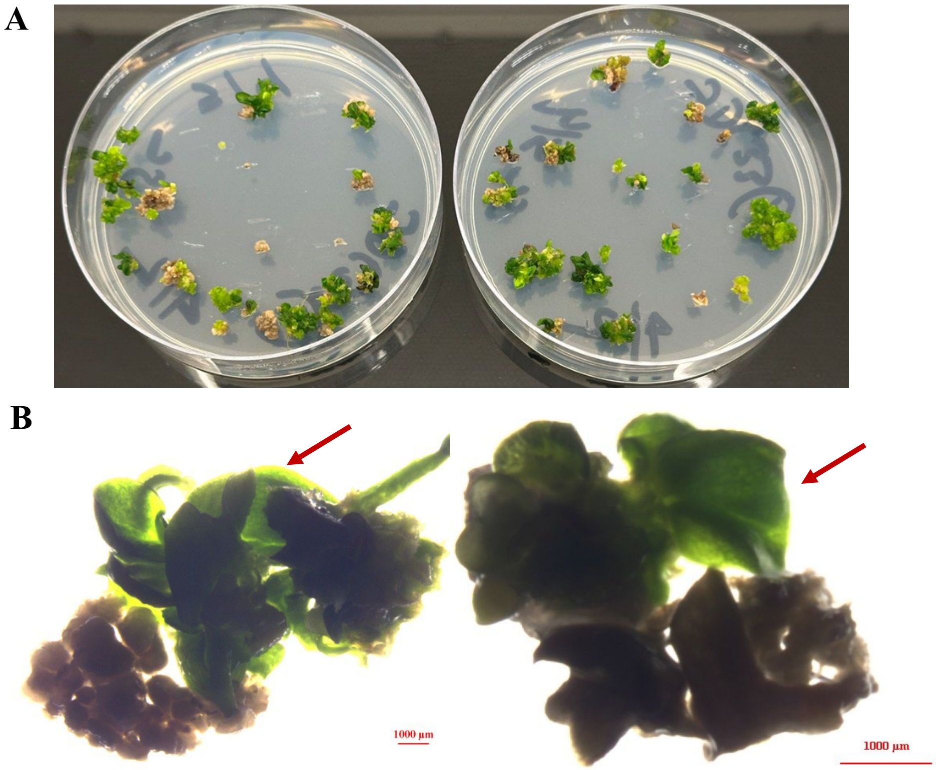


*Note*. (A) Regeneration plates showing putative Cas9-transformed fronds with robust shoot development after 21 days of selection on hygromycin. (B) Red arrows indicate shoot formation from Cas9 positive calli

**Figure S16. Confirmation of gRNA Cloning into the JD633 Plasmid via PCR Analysis**

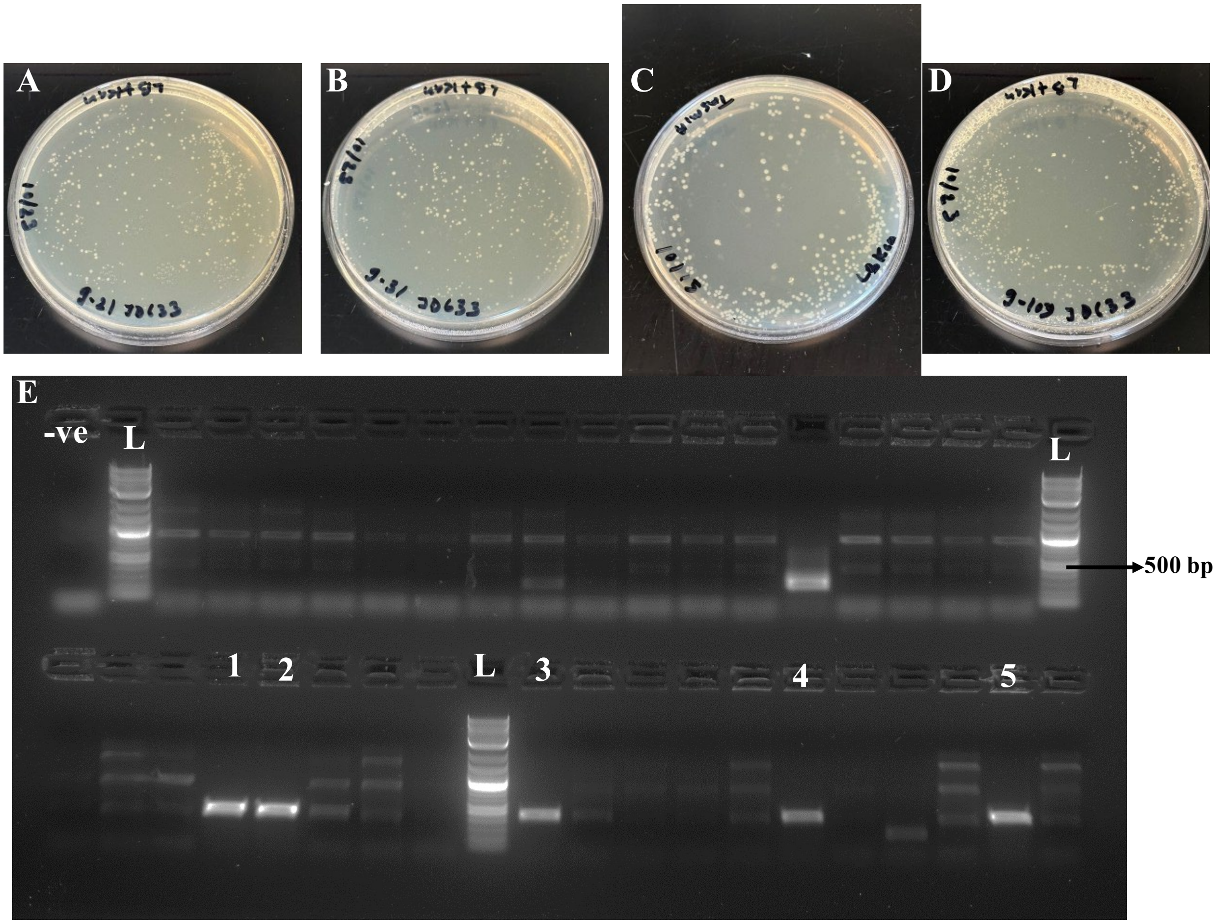


Note. (A–D) LB agar plates showing bacterial colonies following cloning and transformation of individual gRNAs. (E) Agarose gel electrophoresis results of colony PCR using gRNA-specific primers. 1 $kb^+$ DNA ladder was used for size estimation; expected band size (500 bp) indicated. Lanes 1-2: gRNA-90; Lane 3: gRNA-21; Lane 4: gRNA-31; Lane 5: gRNA-109.

**Figure S17. Additional Microscopic Images of Regenerated Fronds of *S. polyrhiza* Containing g-90RNACas9 Construct**

A

B

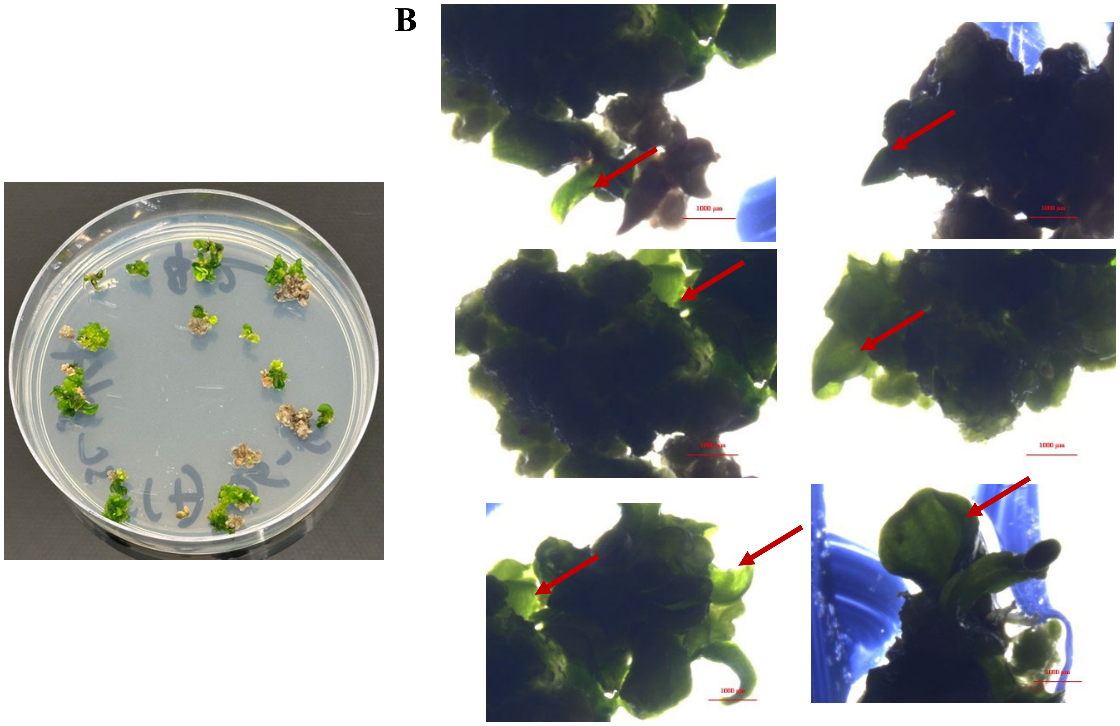

*Note*. (A) Plates containing putative g-90RNACas9-transformed fronds show visible shoot regeneration, indicating survival under selection pressure. (B) Red arrows indicate shoot formation from g-90RNACas9 positive calli.

**Figure S18. PCR Amplification of gRNA-90Cas9 Transgenic *S. polyrhiza* Lines Using PDS1-Specific Primers**

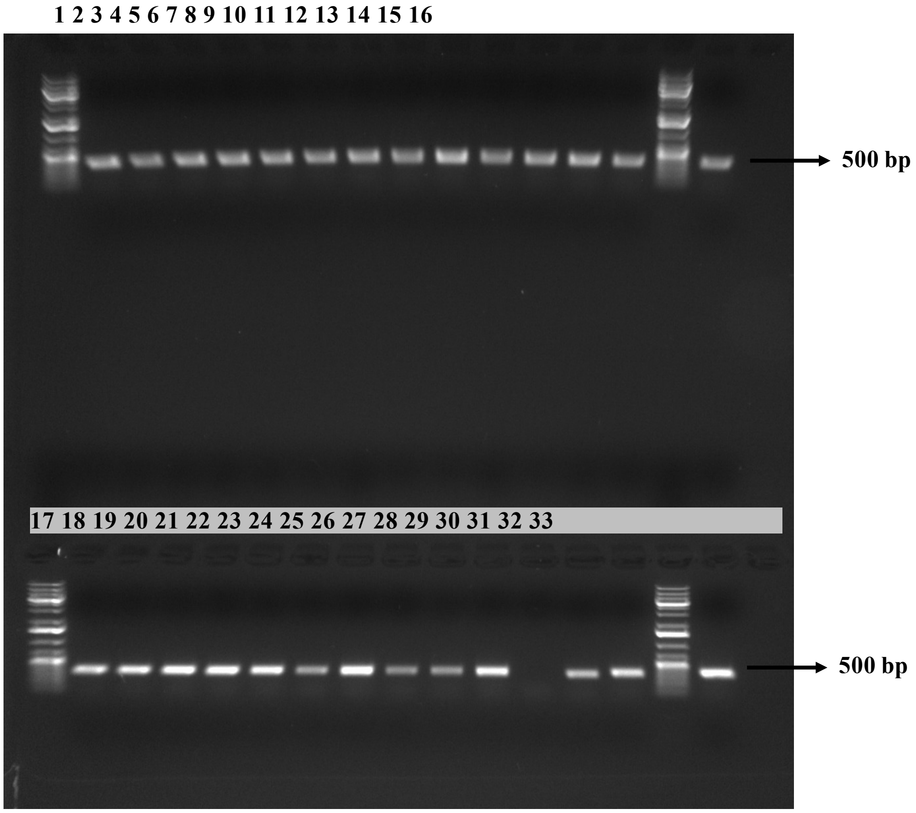


*Note.* Agarose gel electrophoresis shows PCR amplification using primers specific to the PDS1 gene (~348 bp). Lanes 1, 15, 17, and 31 contain the 1 kb$^+$ DNA ladder with the 500 bp band indicated for reference. Lanes 2 and 18 represent wild-type (WT) and control 2 samples, respectively, which show no amplification, confirming the absence of transgene-specific sequences. Lanes 3–14 and 19–30 correspond to gRNA-90Cas9 transgenic lines, all of which show amplification with the PDS1-specific primers. Lanes 16 and 32 represent Cas9 transgenic lines 1 and 2, both showing amplification with the same primer set.

Table S1. guide RNAs Used for Cloning

| | |
|---|---|
| TI-21f-T | TACTTTCGTCTGCGTGGACTATCCG |
| TI-21f-B | AAACCGGATAGTCCACGCAGACGA |
| TI-90r-T | ACTTTCCAATGGTCTAGCAGGACG |
| TI-90r- | AAACCGTCCTGCTAGACCATTGGA |
| TI-31f-T | ACTTGGACTATCCGAGGCCAGAAT |
| TI-31f-B | AAACATTCTGGCCTCGGATAGTC |
| TI-109f-T | ACTTTCCGCGTCCTGCTAGACCAT |
| TI-109f-B | AAACATGGTCTAGCAGGACGCGGA |

Table S2. Primer Used in Molecular Confirmation

| | |
|---|---|
| Primer 1 | 5′-TGGTCGACGTGTTCACGATT-3′ |
| Primer 2 | 5′-AAGGCGGGAAACGACAATCT-3′ |
| Primer 3 | 5′GGTACCATGCTGAAGGAGCTCTACTAC-3′ |
| Primer 4 | 5′-CTCACATACTCCCTGCAGCAGGGTAAG-3′ |
| Primer 5 | 5′-AAGGACTGGGACCCAAAGAAGTAC-3′ |
| Primer 6 | 5′-CGTCAGGGGTGAAGAGATGGATGAT-3′ |
| Primer 7 | 5′-CACGGCGGCGCAGTACGAGGAGC-3′ |
| Primer 8 | 5′-GAAGACGCCGGCGTTCATCATGCTG-3′ |
| Primer 9 | 5′-ATGAAGATGACCTACCACATGGAC -3 |
| Primer 10 | 5′- CTTCTTGGTCTTGTAGGAGGTGTG -3′ |
| Primer 11 | 5′- CTTATGCCCAGCATCCGCTA-3 |
| Primer 12 | 5′- GTCTGCGTGGACTATCCGAG-3′ |
